# Childhood cancer mutagenesis caused by a domesticated DNA transposase

**DOI:** 10.1101/2022.07.05.498128

**Authors:** Ross Keller, Makiko Yamada, Daniel Cameron, Hiromichi Suzuki, Reeti Sanghrajka, Jake Vaynshteyn, Jeffrey Gerwin, Francesco Maura, William Hooper, Minita Shah, Nicolas Robine, Philip Demarest, N. Sumru Bayin, Luz Jubierre, Casie Reed, Michael D. Taylor, Alexandra L. Joyner, G. Praveen Raju, Alex Kentsis

## Abstract

Genomic rearrangements are a hallmark of most solid tumors, including medulloblastoma, one of the most common brain tumors in children. Childhood cancers involve dysregulated cell development, but their mutational causes remain largely unknown. One of the most common forms of medulloblastoma is caused by ectopic activation of Sonic Hedgehog (SHH) signaling in cerebellar granule cell progenitors, associated with genetic deletions, amplifications, and other oncogenic chromosomal rearrangements. Here, we show that *PiggyBac Transposable Element Derived 5 (Pgbd5)* promotes tumor development in multiple developmentally-accurate mouse models of SHH medulloblastoma. Most mice with *Pgbd5* deficiency do not develop tumors, while *Pgbd5*-deficient mice maintain largely normal cerebellar development. Mouse medulloblastomas expressing *Pgbd5* exhibit significantly increased numbers of somatic structural DNA rearrangements, with PGBD5-specific transposon sequences at their breakpoints. Similar sequence breakpoints recurrently affect somatic DNA rearrangements of known tumor suppressors and oncogenes in medulloblastomas in 329 children. Therefore, this study identifies PGBD5 as a primary medulloblastoma mutator and provides a genetic mechanism responsible for the generation of somatic oncogenic DNA rearrangements in childhood cancer.

**One-Sentence Summary:** Induction of somatic oncogenic mutations by the DNA transposase PGBD5 in cerebellar progenitor cells promotes medulloblastoma development.

---

Cancer development is caused by the acquisition of somatic mutations affecting tumor suppressors and oncogenes that encode factors regulating cell proliferation, differentiation, and other hallmarks of cancer (*1*). Compared to aging-associated cancers, childhood tumors are characterized by significantly lower total numbers of genetic mutations, although they exhibit chromosomal deletions, amplifications, translocations, and other complex oncogenic genomic rearrangements (*2*). The origins of these oncogenic mutations in childhood and young adult tumors remain obscure. A quintessential example is medulloblastoma, one of the most frequent childhood brain tumors. For example, medulloblastomas with activation of Sonic Hedgehog (SHH) signaling are caused by the aberrant proliferation of cerebellar granule cell precursors (GCPs) due to activating mutations in *SMO* and somatic deletions of the tumor suppressors *PTCH1, SUFU,* and *TP53*. Medulloblastomas also exhibit frequent amplifications of oncogenes such as *GLI2*, *MYC,* and *MYCN*, and other complex genomic rearrangements. Medulloblastomas can be caused by Li-Fraumeni syndrome and germline deficiency of *TP53*, which dysregulates chromosome replication and repair, thereby causing complex oncogenic DNA rearrangements known as chromothripsis. However, the causes of somatic DNA rearrangements in sporadic medulloblastomas are not known, despite being a defining feature of this and most other childhood and young adult tumors.

Recently, we found that most human medulloblastomas express *PGBD5*, the most evolutionarily conserved transposable element derived nuclease gene in vertebrates, which retains DNA transposition activity in human cells (*3*). We hypothesized that dysregulation of PGBD5 nuclease activity might contribute to the somatic induction of oncogenic DNA rearrangements. Among the four major types of medulloblastomas, *PGBD5* gene expression is highest in tumors with constitutive SHH signaling (*3*). Thus, we used mouse models of sporadic SHH medulloblastoma, induced by constitutive activation of SHH signaling in developing mouse cerebellar GCPs (Fig. 1A).

**Fig. 1.**
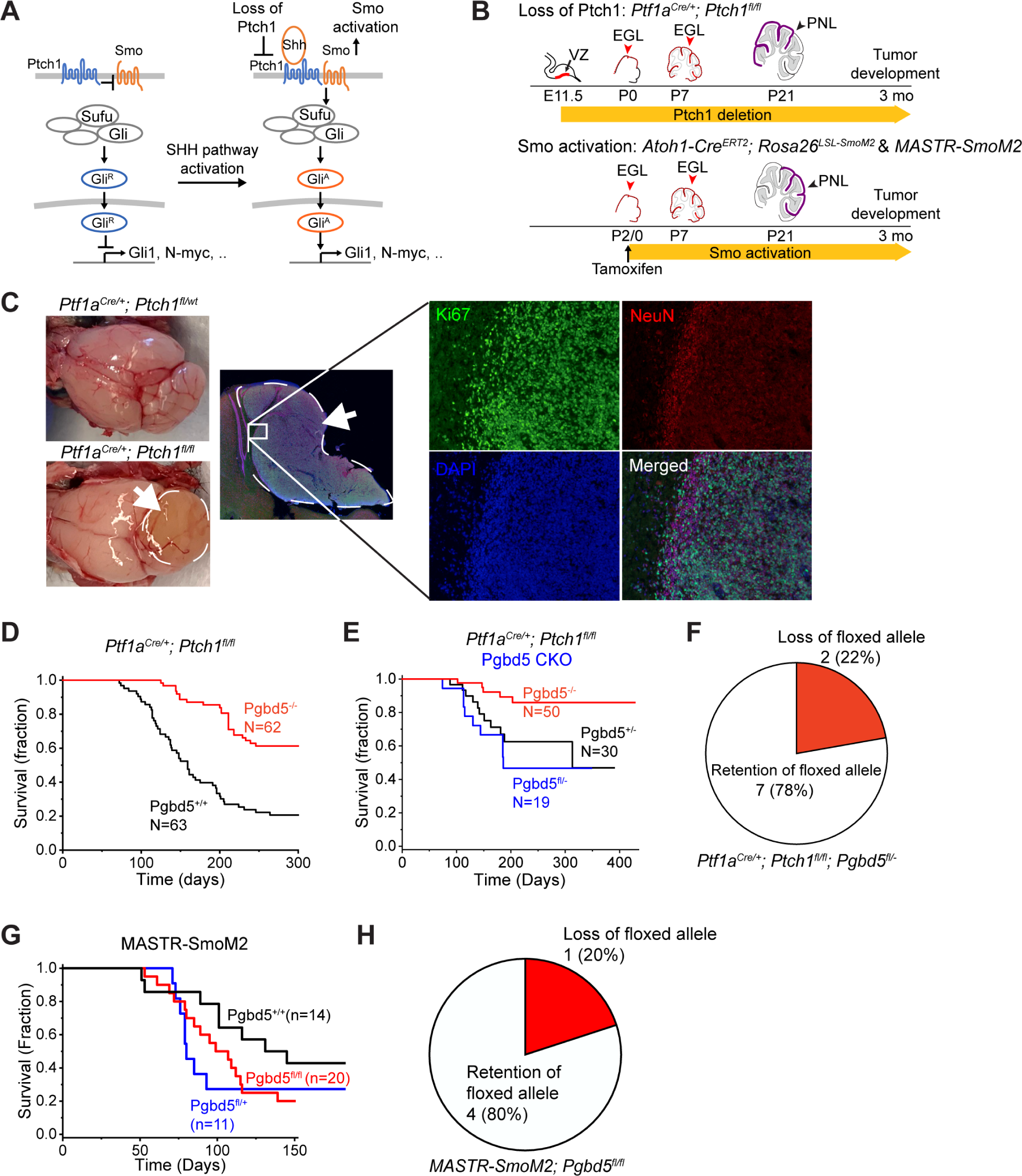
*Pgbd5* promotes tumorigenesis in diverse developmentally accurate mouse models of SHH medulloblastoma. (**A**) Schematic of aberrant mechanisms of SHH signaling in cerebellar granule cell precursors in medulloblastoma (MB) development (left). In *Ptch1-*mutant *Pft1a^Cre/+^;Ptch1^fl/fl^*MB, by deletion of *Ptch1* encoding a receptor for SHH, SMO signaling is disinhibited and highly activated, leading to the generation of activated GLI (GLI^A^). In *Smo*-mutant *MASTR-SmoM2* or *Atoh1-Cre^ERT2^;Rosa26^LSL-SmoM2^* MB, oncogenic constitutively activated form of SMO results in GLI activation and aberrant SHH signaling. (**B**) Schematic of cerebellar tumor development in *Ptch1*- (top) and *SmoM2*-mutant (bottom) MB. Red arrowheads mark conditionally gene-targeted cell populations. Preneoplastic lesions (PNLs, purple) lead to MB development. E and P denote embryonic and postnatal days, respectively, of *Ptch1* deletion (top) and mutant *SmoM2* expression (bottom) upon tamoxifen administration. (**C**) Representative photographs of dissected brains of *Ptf1a^Cre/+^;Ptch1^fl/fl^*(bottom left) mice which develop MBs marked by white arrows and dashed circles, as compared to *Ptf1a^Cre/+^;Ptch1^fl/wt^* mice (top left) which do not develop tumors. Immunofluorescence microscopy (right) shows high Ki67 (green) and low NeuN (red) expression in MB tumors, with nuclei marked with DAPI (blue). The edge of the tumor (white inset) is magnified with NeuN-positive cells on tumor edge corresponding to normal cerebellum. (**D**) Survival of *Ptf1a^Cre/+^; Ptch1^fl/fl^* mice (red) shows that 61% of *Pgbd5^-/-^* mice are tumor-free after 10 months whereas most *Pgbd5^+/+^* mice (black) develop tumors (*n* = 50), as analyzed in 3 independent cohorts (Fig. S1B). (**E**) Survival of *Pft1a^Cre/+^;Ptch1^fl/fl^*mice with conditional KO (CKO) of *Pgbd5^fl/-^* (blue) or control *Pgbd5^+/-^* (black) or *Pgbd5^-/-^* (red) littermates. (**F**) Genomic PCR analysis of conditional *Pgbd5* excision in *Pgbd5^fl/-^* CKO *Ptch1*-mutant tumors demonstrates that 7 out of 9 (79%) analyzed tumors retain intact *Pgbd5*, as detailed in Fig. S1B. (**G**) Survival of *MASTR-SmoM2* mice with conditional knockout of *Pgbd5^fl/fl^*(red), as compared to control *Pgbd5^+/+^* (black) or *Pgbd5^fl/+^*(blue) littermates. (**H**) Genomic PCR analysis of conditional *Pgbd5* excision in *MASTR-SmoM2; Pgbd5^fl/fl^* tumors demonstrates that 4 out of 5 (80%) analyzed tumors retain *Pgbd5* floxed alleles, as detailed in Fig. S2B.

First, we engineered mice with *loxP* sites flanking exon 4 of mouse *Pgbd5*, generating floxed *Pgbd5^fl/fl^* and knockout *Pgbd5^-/-^* mice in which translation of the protein is out of frame after exon 4 (Fig. S1A). In situ hybridization with a *Pgbd5* exon 4-specific probe set confirmed complete loss of exon 4 transcripts in *Pgbd5^-/-^* mice (Fig. S1B & S3B). We then crossed the *Pgbd5^-/-^* alleles into *Ptf1a^Cre/+^;Ptch1^fl/fl^* mice, which develop tumors with the highest known penetrance among mouse models of sporadic SHH medulloblastoma (Fig. 1B & C) (*4*). Mosaic loss of *Ptch1* leads to pre-neoplastic hyperplasia of developing cerebellar GCPs due to the constitutive activation of SHH signaling, similar to sporadic human SHH medulloblastomas with somatic mutations of *PTCH1*, which affect more than 40% of patients with SHH medulloblastomas (*5*). Analysis of three independent cohorts of *Pgbd5^wt/wt^;Ptf1a^Cre/+^;Ptch1^fl/fl^*and *Pgbd5^-/-^;Ptf1a^Cre/+^;Ptch1^fl/fl^* mice demonstrated a significant and pronounced requirement of *Pgbd5* for medulloblastoma development (Fig. 1D & S1C). Among *Pgbd5*-knockout tumor model mice, as many as 70% of animals did not develop tumors after one year of life (mean 61%, log-rank *p* = 1.4e-8), whereas the majority of *Pgbd5^wt/wt^* mice (79%) rapidly succumbed to medulloblastomas with a median latency of five months.

To exclude the possibility of tumor cell extrinsic effects of germline *Pgbd5^-/-^* deletion, we used *Pgbd5*-floxed mice in which *Pgbd5* loss was primarily confined to cerebellar progenitor cells due to conditional Cre expression and *loxP* recombination. Both *Pgbd5^fl/-^;Ptf1a^Cre/+^;Ptch1^fl/fl^* mice and their *Pgbd5^wt/-^;Ptf1a^Cre/+^;Ptch1^fl/fl^*littermates developed medulloblastomas with similar penetrance and latency (Fig. 1E). However, genomic PCR analysis using primers specific for the *Pgbd5*-floxed allele demonstrated that 7 out of 9 analyzed tumors (78%) retained a substantial amount of intact *Pgbd5*, indicating a selective advantage for *Pgbd5*-expressing tumor progenitor cells (Fisher’s exact test *p* = 2.3e-3; Fig. 1F & S1C). We performed a similar analysis using a developmentally accurate mouse model of sporadic *SMO*-mutant medulloblastoma, corresponding to a mutation of *SMO* that aberrantly activates SHH signaling in human patients (*5, 6*).

This model leverages a system for mosaic mutagenesis with spatial and temporal control of recombination (MASTR) (*7, 8*). In this system, GFP-Cre is induced by tamoxifen at postnatal day 0 in cerebellar granule cell precursors, and further induces oncogenic *SmoM2* expression and concurrent deletion of *Pgbd5* (Fig. S2A). Both *Pgbd5^fl/fl^* and their *Pgbd5^fl/+^* littermates developed tumors with similar latencies (Fig. 1G). However, genomic PCR analysis again showed that the majority of analyzed *Pgbd5^fl/fl^* tumors (4 out of 5) retained intact *Pgbd5* alleles, indicating that *Pgbd5* expression enhances *SmoM2*-mutant SHH medulloblastoma development (Fisher’s exact test *p* = 2.1e-2; Fig. 1H & S2B). To exclude the possibility that apparent *Pgbd5* expression was due to infiltration of stromal or immune cells, we confirmed the requirement for *Pgbd5* expression in MB tumor cell development using three of the same *SmoM2*-mutant tumors by *in situ* hybridization with a *Pgbd5* exon 4-specific probe and *Cre*-specific probe as a positive control for tumor cells (Fig. S3C-F). This revealed specific *Pgbd5* transcript expression in tumor cells 2 out of 3 of analyzed *MASTR-SmoM2;Pgbd5^fl/fl^* tumors. In all, these results indicate that *Pgbd5* promotes tumor development in diverse developmentally accurate mouse models of SHH medulloblastomas.

SHH medulloblastomas originate in developing cerebellar GCPs, which in turn are dependent on physiologic SHH signaling (*7, 9–13*). To exclude the possibility that Pgbd5-induced tumorigenesis is due to its control of normal cerebellar development, we analyzed the cerebellar cytoarchitecture of *Pgbd5^-/-^* mice. We observed grossly intact medial cerebellar vermis and lateral hemispheres, including normal cytoarchitecture and morphology, which are essential hallmarks of cerebellar development (Fig. 2A). To examine the effects of *Pgbd5* deficiency on SHH signaling directly, we isolated GCPs from the cerebella of 5-day old mice, when SHH signaling is required for cerebellar development, and measured SHH pathway activity using quantitative RT-PCR of the canonical SHH signaling biomarker *Gli1* (*11*). We observed no significant differences in *Gli1* expression between *Pgbd5^-/-^* and wildtype developing cerebellar GCPs (*p* = 0.87; Fig. 2B). Thus, the requirement of *Pgbd5* for SHH medulloblastoma development cannot be explained by the effects of SHH signaling on growth and survival of mutant GCPs which initiate tumor progression.

**Fig. 2.**
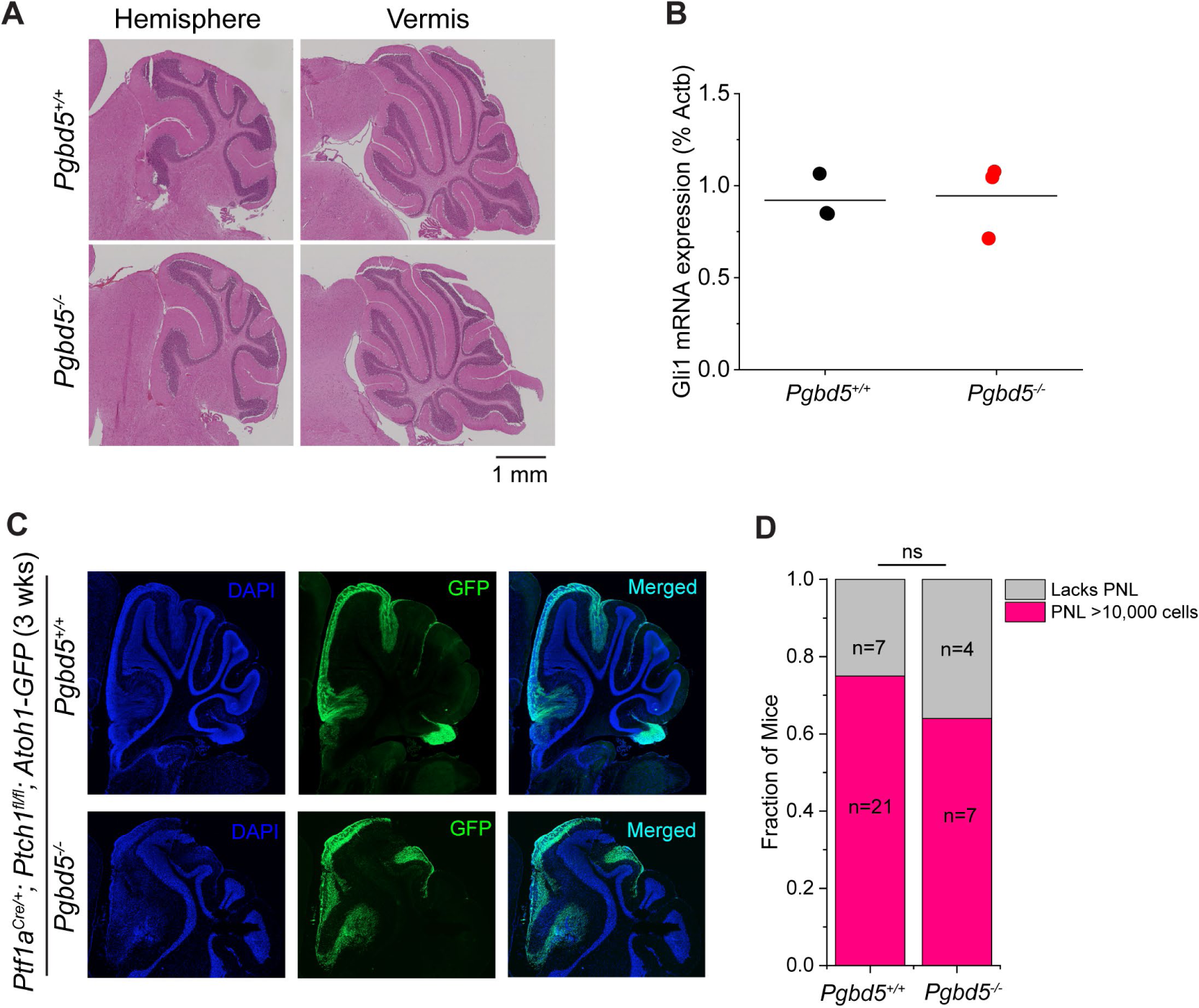
*Pgbd5* is dispensable for normal SHH signaling and cerebellar development. (**A**) Representative immunohistochemistry micrographs of sagittal sections of cerebellum of *Pgbd5^+/+^*(top) and *Pgbd5^-/-^* (bottom) mice at 6 weeks of age show normal cytoarchitecture and morphology of cerebellar hemispheres (left) and vermis (right). (**B**) Expression of *Gli1* mRNA in purified cerebellar granule cell precursors from 5-day old *Pgbd5^+/+^* (black) and *Pgbd5^-/-^* (red) mice. Bars represent means of three biologic replicates (*p* = 0.87). (**C**) Representative fluorescence images of cerebellar hemispheres of 3-week-old *Pgbd5^+/+^* (top) versus *Pgbd5^-/-^* (bottom) *Ptf1a^Cre/+^; Ptch1^fl/fl^;Atoh1-GFP* mice showing preneoplastic lesions (green) marked by Atoh1-GFP expression, with nuclei marked with DAPI (blue). (**D**) Fraction of mice harboring preneoplastic lesions (PNLs in red) in *Pgbd5^wt/wt^;Ptf1a^Cre/+^;Ptch1^fl/fl^;Atoh1-GFP* and *Pgbd5^-/-^;Ptf1a^Cre/+^;Ptch1^fl/fl^;Atoh1-GFP* mice between 3 to 8 weeks of age. Both groups harbor similar fractions of PNLs which are defined as at least 10,000 Atoh1-GFP-positive cells (Fisher’s exact test *p* = 0.69); ns, not significant.

Oncogenic SHH signaling leads to hyperplasia of cerebellar GCPs. These pre-neoplastic cells express Atoh1 and persist in the external granule layer between 3-8 weeks of age (*14, 15*). To examine the effects of Pgbd5 on pre-neoplastic GCP hyperplasia, we used an *Atoh1-GFP* reporter transgene and fluorescence-activated cell sorting (FACS) to isolate pre-neoplastic GCPs developing in the cerebellar external granule layer in 3 to 8-week old *Pgbd5^-/-^;Ptf1a^Cre/+^;Ptch1^fl/fl^;Atoh1-GFP* mice (*16*). We found that 21 out of 28 (75%) of *Pgbd5^wt/wt^;Ptf1a^Cre/+^;Ptch1^fl/fl^;Atoh1-GFP* mice harbored pre-neoplastic cells in their cerebella, which was similar to *Pgbd5*-knockout mice (Fig. 2C-D). The pre-neoplastic populations also showed no significant differences in the numbers of *Atoh1*-expressing cells (*p* = 0.16; Fig. S4). Thus, *Pgbd5* is dispensable for normal cerebellar development, physiologic SHH signaling, and the growth of pre-neoplastic SHH medulloblastoma progenitor cells.

Vertebrate *PGBD5* is derived from *piggyBac* DNA transposases which use an RNAse H-like nuclease domain to catalyze DNA double-strand breaks and rearrangements at specific sequences (*3, 17–19*). Does PGBD5 promote medulloblastoma development by inducing sequence-specific somatic mutations? To test this hypothesis, we performed whole-genome PCR-free DNA sequencing of medulloblastomas from both *Ptch1*- and *SmoM2*-mutant tumors, as compared to their matched normal tissues. We analyzed resultant sequencing data using recently developed methods optimized for the accurate detection of somatic cancer genome rearrangements (*20*).

*Pgbd5*-expressing medulloblastomas exhibited nearly twice as many somatic DNA rearrangements, including deletions, insertions, translocations, and amplifications, as compared to the relatively few tumors that could be isolated from *Pgbd5*-deficient mice (mean 69 and 44, respectively; t-test *p* = 0.036; Fig. 3A). This increase could not be attributed to the age of tumor development, as *Pgbd5*-expressing medulloblastomas were significantly younger (mean 140 and 201 days, respectively; t-test *p* = 0.024; Fig. 3B). Indeed, there was no correlation between the number of somatic genome rearrangements and tumor age (*r* = −0.18, *p* = 0.55; Fig. S5A), consistent with a distinct somatic mutational process responsible for tumor development. Similar to human medulloblastomas, both *Ptch1*- and *SmoM2*-mutant mouse medulloblastomas exhibited relatively low numbers of single nucleotide variants, consistent with their early embryonal age of onset, regardless of *Pgbd5* expression (mean 1.6 and 0.28 mutations/megabase, respectively; Fig. S6A-C). There were no significant differences in single- or double-nucleotide mutational signatures between *Pgbd5^-/-^* and *Pgbd5^+/+^*tumors, with predominance of SBS5 and SBS18 signatures due to chronological age and radical oxygen stress damage, respectively (Fig. S6C) (*21*).

**Fig. 3.**
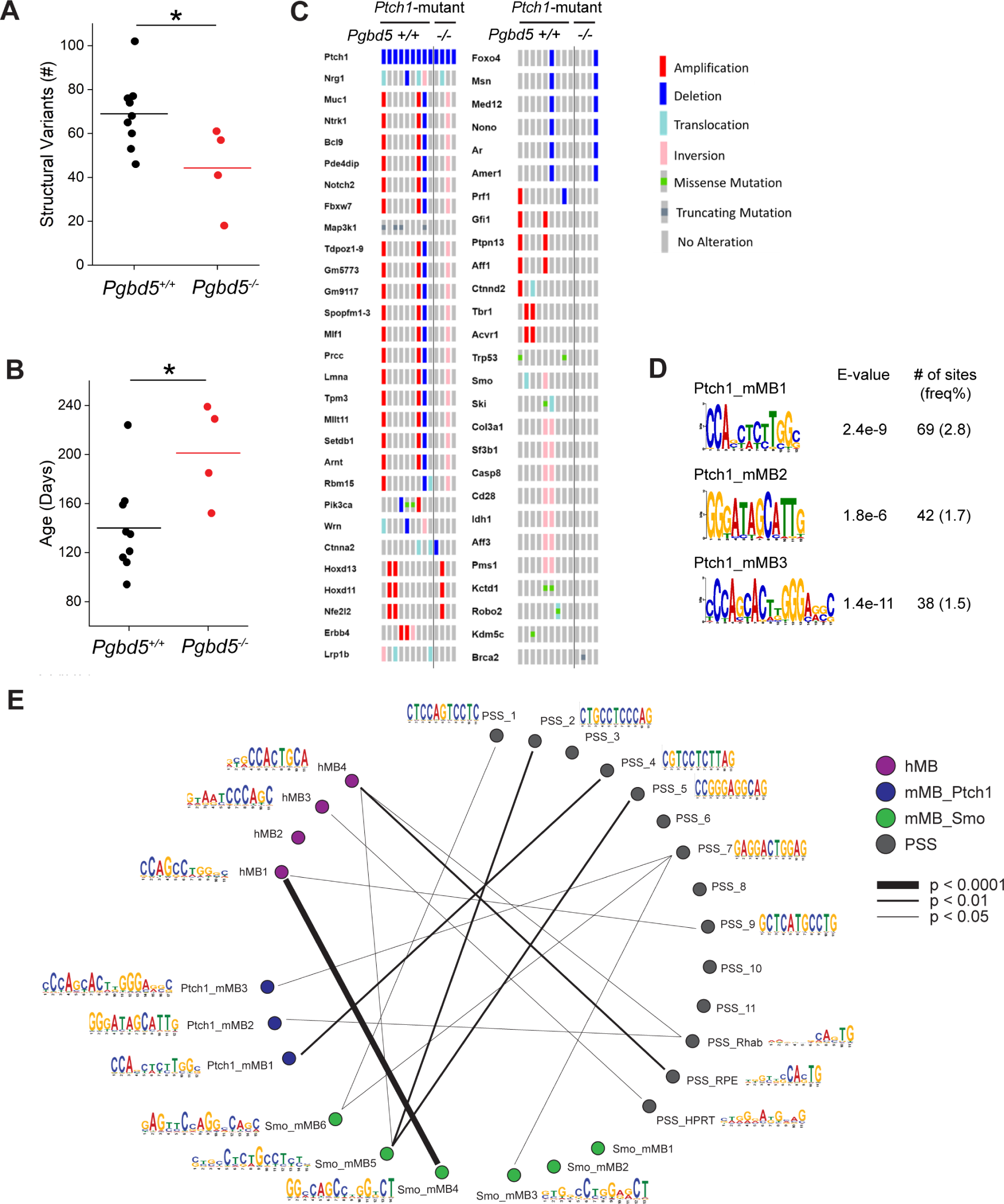
*Pgbd5* promotes somatic mutagenesis of recurrently mutated tumor suppressor and oncogenes in mouse SHH medulloblastomas. (**A**) Numbers of structural variants in *Ptf1a^Cre/+^; Ptch1^fl/fl^* tumors. *Pgbd5^+/+^* tumors (black, *n* = 9) harbor more structural variants than *Pgbd5^-/-^* (red, *n* = 4). Lines indicate mean (69 and 44, respectively) and significance is measured using t-test (\**p* = 0.036). (**B**) The age of tumors (days) in *Ptch1*-mutant tumors. *Pgbd5^+/+^* tumors (black, *n* = 9) are younger than *Pgbd5^-/-^* tumors (red, *n* = 4; mean 140 and 201 days, \**p* = 0.024). (**C**) Oncoprint showing genes recurrently affected by structural variants and SNVs in independent *Ptch1*-mutant tumors. Genes are curated based on likelihood that SVs or SNVs affect gene function (see methods for details). The left nine and right four columns indicate tumors from *Pgbd5^+/+^* and *Pgbd5^-/-^* mice, respectively. Red, blue, light blue, pink, and gray symbols indicate amplifications, deletions, translocations, inversions, and no alteration, respectively. Green and dark gray squares in gray symbols indicate missense and truncating mutations, respectively. (**D**) Three *Pgbd5^+/+^*-specific motifs are identified at structural variant breakpoints in *Ptch1*-mutant tumors, using discriminative MEME with *Pgbd5^-/-^* tumors as controls. E-values indicate MEME discriminative algorithm significance (see methods for details). The frequency shown was calculated by dividing the number of sites by total numbers of 50bp breakpoint sequences extracted from structural variants. (**E**) Circos plot showing similarities among all motifs. Line thickness is proportional to degree of similarity, as indicated. The mouse MB motifs include three *Ptch1*-mutant motifs in blue (Ptch1_mMB1-3) and six *SmoM2*-mutant motifs in green (Smo_mMB1-6). The human MB motifs are colored purple (hMB1-4). The previously identified PGBD5-specific signal sequence (PSS) motifs include 14 motifs in gray (PSS_1-11, PSS_Rhab, PSS_RPE, and PSS_HPRT) (*3, 19*).

We identified putative MB tumor suppressor and oncogenes arising from *Pgbd5*-induced genomic rearrangements by analyzing their recurrence in independent mouse tumors, as compared to genes recurrently mutated in human medulloblastomas (Fig. 3C & S7, and Data S1-2, & S5). This identified several genes, including *Fbxw7*, *Tbr1*, *Gfi1*, *Pik3ca,* and others known to be recurrently mutated in human SHH and non-SHH medulloblastomas (Data S8) (*5*). Importantly, the same genes were also recurrently mutated in *SmoM2-*mutant mouse MB tumors, but not in the rare *Pgbd5*-deficient tumors (Fig. S8A). This suggests that mouse SHH tumors model salient mutational features of human medulloblastomas, including specific developmental PGBD5-induced mutations.

To elucidate specific mutational processes responsible for PGBD5-induced somatic genomic rearrangements in SHH medulloblastomas, we extracted 50-basepair sequences flanking all somatic DNA rearrangement breakpoints and analyzed their composition using supervised and de novo sequence motif analysis algorithms (Fig. S9A-B, Data S6). This analysis showed that most of the somatic DNA rearrangements contained repetitive sequences at their breakpoints, consistent with involvement of non-allelic homologous recombination (NAHR) or microhomology-mediated end-joining (MMEJ), which showed modest but not significant differences between *Pgbd5*-expressing and *Pgbd5*-deficient SHH medulloblastomas (Fig. S10A).

In contrast, specific non-repetitive sequence breakpoints were significantly enriched at somatic DNA rearrangements in *Pgbd5*-expressing SHH mouse medulloblastomas as compared to the relatively few *Pgbd5*-deficient tumors (mean 18 versus 6 per tumor, respectively; t-test *p* = 2.7e-3; Fig. S10A-B). Importantly, of the 2480 breakpoints of somatic DNA rearrangements among nine *Ptch1*-mutant *Pgbd5^+/+^* medulloblastoma tumors, 149 exhibited distinct sequence motifs (Fig. 3D). These breakpoint sequences exhibited significant similarity to the PGBD5-specific signal sequence (PSS) motifs previously observed to be rearranged by PGBD5 in genomic transposition and forward genetic assays (*p* = 7.6e-3, 4.5e-2, and 2.4e-2, respectively; Fig. 3E & Table S1) (*3, 19*). We confirmed the specificity of this PSS breakpoint detection using shuffled sequences which showed no significant association in spite of having identical sequence composition (Fig. S12 & Table S1). We observed similar results in *SmoM2*-mutant MB tumors (252 out of 5036 PSS-like breakpoints; *p* = 5.2e-3 – 3.4e-2; Fig. S11, 3E & S12, and Table S1). The relatively modest but specific association of MB genomic rearrangement breakpoints with previously observed PSS motifs suggests that additional sequences of PGBD5 and other sequence-specific developmental mutational processes remain to be discovered. These findings indicate that PGBD5-induced sequence-specific somatic mutagenesis contributes to mouse SHH medulloblastoma development.

To examine the contribution of PGBD5-induced sequence-specific somatic mutagenesis in human medulloblastomas, we used de novo local sequence assembly-based methods to analyze 329 tumor genomes isolated from patients with all four major MB tumor subtypes, including SHH medulloblastomas (Fig. 4A-B) (*22*). We found that nearly one in five somatic medulloblastoma DNA rearrangement breakpoints contained PSS motifs, previously observed to be rearranged by PGBD5 in genomic transposition and forward genetic assays (Fig. 4C, Data S9-12). This enrichment was significant when compared to somatic breakpoints in non-PGBD5-expressing but highly somatically rearranged human breast carcinomas (χ^2^ test *p* = 7.7e-117; Fig. 4C) (*3, 23*). Using unsupervised de novo sequence motif analysis, we also identified four sequence motifs, termed hMB1-4, which were similarly specifically and significantly enriched in breakpoint sequences of somatic human DNA medulloblastoma rearrangements, but not human breast carcinomas (χ^2^ test *p* = 1.5e-155; Fig. 4D).

**Fig. 4.**
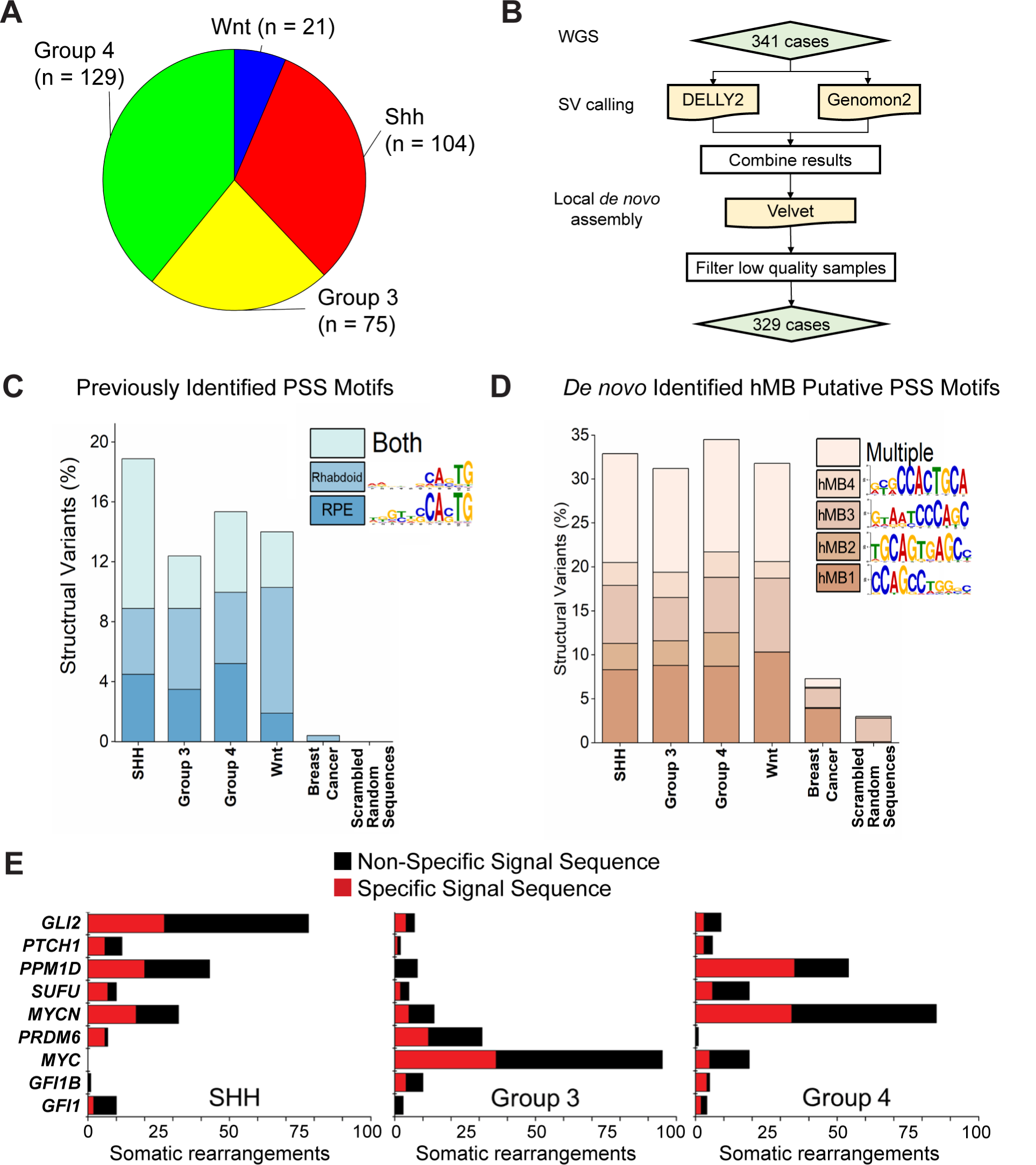
PGBD5-associated sequence breakpoints recurrently affect somatic DNA rearrangements of known tumor suppressor and oncogenes in human medulloblastomas. (**A**) Representation of human MB patient cohort showing the four major subgroups of medulloblastoma that were included in the analysis (*n* = 329). (**B**) Pipeline to identify somatic structural variants in human medulloblastoma (see methods for details). (**C**) Previously identified PSS sequences (*3, 19*) are enriched at structural variant breakpoints in human medulloblastoma as compared to somatic structural variants in human breast carcinomas (χ^2^ test *p* = 7.7e-117). Percentages represent the frequency of structural variants (each SV has two breakpoints and 4×50mers) where the motif was identified using a FIMO q-value threshold of 0.3 based on a receiver operating characteristic curve analysis (Fig. S13B & C). Multiple indicates more than one motif was identified at one structural variant, either in distinct or the same 50mer. Scrambled sequences showed no enrichment and represent the background of the FIMO algorithm. (**D**) A set of four de novo motifs identified at SV breakpoints in human medulloblastoma are enriched relative to breast carcinoma and scrambled sequences. hMB1-4 were identified as being specific using MEME and eliminating repetitive motifs. Additionally, discriminative MEME, where control sequences were a set of 50,000 randomly selected 50mers from the hg19 reference genome, was used to determine whether the motif was enriched at breakpoints relative to the genome (Fig. S13). Percentages represent motif frequency among structural variants as in (C) and are compared to structural variant breakpoints in human breast carcinoma (χ^2^ test *p* = 1.5e-155) and scrambled sequences, which represent the background of the FIMO algorithm. (**E**) Recurrently mutated MB tumor suppressors and oncogenes in diverse tumor subtypes involve somatic DNA rearrangements with specific (red) sequence breakpoints, including PSS motifs. Numbers refer to the structural variants detected in human patient cohort described in (A).

In total, nearly one in three somatic human medulloblastoma DNA rearrangements exhibited specific sequence breakpoints (Fig. 4D). Multiple human tumor DNA rearrangements breakpoints were similar to those detected in Pgbd5-induced mouse SHH medulloblastomas (*p* = 5.3e-6 and 1.0e-2 for hMB1 versus Smo_mMB4 and hMB4 versus Smo_mMB5, respectively; Fig. 3E & S12 and Table S2). In particular, hMB4 motif showed significant similarity to PSS sequences identified as direct PGBD5 substrates (*p* = 3.3e-2 and 6.0e-4 for PSS_Rhab and PSS_RPE, respectively; Fig. 3E & S12 and Table S1). Consistent with the oncogenic activity of PGBD5-induced somatic mutagenesis, many recurrently mutated key SHH and non-SHH medulloblastoma tumor suppressors and oncogenes, including *GLI2*, *PPM1D*, and *MYC* (*24, 25*), involved breakpoints with specific PSS-like sequences (Fig. 4E and Data S5). Therefore, human medulloblastomas are defined by somatic DNA deletions, amplifications and other chromosomal rearrangements, marked by specific breakpoints with similarity to PGBD5-specific signal sequences, which recurrently affect medulloblastoma tumor suppressors and oncogenes.

Although somatic mutational mechanisms have been extensively documented in aging-associated cancers, how oncogenic DNA rearrangements occur upon dysregulated development in childhood and young-onset tumors remained poorly understood. Here, we demonstrate that a domesticated DNA transposase is a major cause of oncogenic DNA rearrangements in medulloblastomas, a common childhood brain tumor (*26*). We provide evidence that oncogenic developmental signaling can not only induce preneoplastic cell expansion but also involves endogenous sequence-specific mutators to generate somatic mutations of tumor suppressor and oncogenes required for malignant cell transformation. Dysregulation of domesticated DNA transposases during cancer development likely increases the probability of oncogenic mutations via rearrangements of substrate transposon sequences by recombining genetic loci into new elements. We also derived first principles of how transposase-mediated somatic genome mutagenesis occurs, providing a foundation for the identification of other sequence-specific mutational processes in other cancers. Incorporation of sequence-specific DNA rearrangements into cancer mutational profiling should improve its diagnosis and ultimate treatment (*27, 28*).

Transposase-mediated somatic mutagenesis offers a genetic mechanism for site-specific DNA rearrangements in developmental cancers, including neuroblastomas, Ewing sarcomas, desmoplastic small round cell tumors, rhabdomyosarcomas, and many other young-onset cancers that express PGBD5. Sequence-specific mutators such as PGBD5 also offer a plausible mechanism for the generation of complex DNA rearrangements typically observed in these tumors, including chromoplexy, and multiple other complex DNA rearrangements with distinct structural features (*29*). Many studies have implicated replication stress as a cause of somatic mutations in cancer, including replication stress induced by high level SHH signaling in cerebellar GCPs (*30*). However, this concept does not explain how sequence-specific mutations, including of tumor suppressors and oncogenes, initially occur. The results presented here offer a likely mechanism by which dysregulated SHH developmental signaling activates PGBD5 mutagenic activity and/or impairs its efficient repair. It is also possible that dysregulation of a domesticated DNA transposase such as PGBD5 can induce not only somatic mutagenesis by virtue of its nuclease activity, but also epigenetic dysregulation via its interactions with chromatin and cellular cofactors.

Although we focused on domesticated DNA transposase PGBD5 in this study, we propose that the principles and implications of somatic developmental mutators revealed herein extend beyond cancer development (*26*). Transposases are among the most ubiquitously present genes in living organisms (*31*), with many transposases domesticated in diverse genetic species and somatic tissues (*32*). One can imagine how domesticated transposases can provide molecular mechanisms for somatic genetic diversification during normal tissue development, and when dysregulated, cause somatic mutations which are increasingly being discovered as causes of diverse sporadic human diseases.

## Acknowledgments

We thank Anton Henssen, Gabriella Casalena, Helen Mueller, Sumiko Takao, Shuyuan Cheng, Michael Kharas, Hao Zhu, Alejandro Gutierrez, and Marc Mansour for helpful suggestions, Mithat Gönen for statistical advice, Wenfei Kang, Eric Rosiek, Katia Manova-Todorova, Victor Morell, the MSK Molecular Cytology, Integrated Genomics and Bioinformatics Core facilities, and the Center for Comparative Medicine and Pathology for technical assistance, and Nina Kentsis for editing help. AK is a Scholar of the Leukemia & Lymphoma Society.

## Funding

National Institutes of Health grants R01 CA214812 (AK), R01 CA192176 (ALJ), and P30 CA008748 (AK, ALJ)

St. Baldrick’s Foundation (AK)

Burroughs Wellcome Fund (AK)

Rita Allen Foundation (AK)

Pershing Square Sohn Cancer Research Alliance and the G. Harold and Leila Y. Mathers Foundation (AK)

Starr Cancer Consortium (AK)

Cookies for Kids’ Cancer (AK, GPR)

MSK Brain Tumor Center (AK, GPR)

MSK Functional Genomics Initiative (ALJ)

## Author contributions

Conceptualization: AK, GPR, ALJ, MDT, RK, MY

Methodology: RK, MY, DC, HS, RS, JV, JG, FM, WH, MS, NR, PD, NSB, LJ, CR

Investigation: RK, MY, DC, HS, RS, JV, JG, FM, WH, MS, NR, PD, NSB, CR

Visualization: MY, RK, FM, HS, JV, NSB

Funding acquisition: AK, GPR, ALJ, MDT

Project administration: AK

Supervision: AK, GPR, ALJ, MDT

Writing – original draft: AK, MY, RK

Writing – review & editing: All authors

## Data and materials availability

Sequencing data available from the NCBI Sequence Read Archive (mouse) and the EGA Archive (human) using respective accession numbers listed in Materials and Methods. All processed data, including specific structural variants and their breakpoint sequences, are openly available via Zenodo. Genetically-engineered mouse strains are available from authors upon request.

## Materials and Methods

### Mice

*Ptf1a-Cre/+*, *Ptch1^fl/fl^*, and *Atoh1-GFP* mice were obtained from Mikio Hoshino (*33*), Brandon Wainwright (*34*), and Jane Johnson (*16*), respectively. All three lines were initially maintained on mixed backgrounds and subsequently backcrossed with C57BL/6J mice to generate C57BL/6J-background mice. *Atoh1-CreER^T2^*(*Math1-CreER^T2^*) (007684) and *R26SmoM2* (005130) mice were obtained from the Jackson Laboratory (*35, 36*). *Atoh1-CreER^T2^* mice were maintained on SW background. *Pgbd5*-floxed mice were generated by targeting exon 4 of mouse *Pgbd5* (InGenious Targeting Laboratory; Fig. S1A). Targeting vector consisting of *LacZ* and *NeoR* cassettes was electroporated in C57BL/6 ES cells and targeted clones were microinjected into Balb/c blastocysts. Resulting chimeras were crossed with C57BL/6 FLP mice to remove the Neo cassette to generate *Pgbd5*-floxed mice, as confirmed using genotyping with SC2 (GAG AGC ACC GTT GGT GCA TAT CAG) and SC4 (AGA GTA TGA GCG GGA GAG GAG CAG) (Fig. S1A). *Pgbd5*-floxed mice were then crossed with *B6.FVB-Tg(EIIa-cre)C5379Lmgd/J (EIIa-Cre)* mice to generate global *Pgbd5*-deficient mice, as confirmed by genotyping with SC2 and SC5 (TTT CTC AGC TGT CCC CAG CAT AGC) primers. *Pgbd5*-deficient mice were backcrossed with C57BL/6J mice for six generations. The MASTR model was adapted from Wojcinski et al and Lao et al (*7, 37*) (Fig. S2A). *Atoh1-FlpoER/+*, *R26^MASTR^ (MA)*, and *R26SmoM2 (SmoM2)* mice were maintained on a mixed background and subsequently outbred into a SW background. Genotyping of *Atoh1-FlpoER/+* and *R26^MASTR^* were done by PCR using following pairs of forward and reverse primers: Fwd GCTCTACTTCATCGCATTCCTTGC, Rev ATTATTTTTGACACCAGACCAAC, Fwd GATATCTCACGTACTGACGG, Rev TGACCAGAGTCATCCTTAGC, respectively. Offspring were obtained by crossing either *Atoh1-FlpoER/+*; *SmoM2/SmoM2; Pgbd5^fl/+^* with *MA/+*; Pgbd5^fl/fl^, *Atoh1-FlpoER/+*; *SmoM2/SmoM2*; *Pgbd5^fl/+^* with *MA/MA*, or *Atoh1-FlpoER/+*; *SmoM2/SmoM2*; *Pgbd5^fl/+^* with *MA/MA*; *Pgbd5^fl/+^*. All experiments were conducted in compliance with protocols approved by the Memorial Sloan Kettering Cancer Center Institutional Animal Care and Use Committee.

### Tumor induction

On postnatal day (P) 2 or 0, pups from *Atoh1-CreER^T2^* x *R26SmoM2* or crossings from the MASTR model were injected with 200 mg/kg Tamoxifen (Sigma T5648) in corn oil subcutaneously. Study endpoints included reduced activity, ataxia and/or domed skulls.

### Histology and immunohistochemistry of mouse cerebellum and tumors

Under deep anesthesia, animals were perfused with intracardiac 0.9% saline followed by 4% paraformaldehyde (PFA)/0.1M phosphate buffer (PB). Brains were dissected, and further fixed in 4% PFA/0.1M PB overnight, and embedded in paraffin blocks. Sagittal sections of cerebella were stained with hematoxylin and eosin. Immunohistochemistry was performed using anti-Ki67 (ab15580, Abcam) and anti-NeuN (A60; EMD Millipore), respectively.

### Copy number analysis

Upon euthanasia and tumor dissection, genomic DNA was extracted from tumors and matched normal tissues (Transnetyx). gDNA real-time PCR was performed using probes for *Pgbd5* wildtype (Pgbd5-1 WT), *Pgbd5*-floxed (Pgbd5-1 FL), and *Pbgd5*-null (Pgbd5-1 EX) alleles, with probes for *jun* as reference (Transnetyx).

### Isolation of mouse cerebellar granule cell precursors (GCPs)

The protocol was adapted from Nakashima et al and Lee et al (*38, 39*). Briefly, P5 cerebella were trypsinized at 37℃, followed by DNaseI treatment. GCPs were then isolated by a Percoll gradient consisting of 60% and 35% Percoll solutions. After centrifugation at 2000g, cells at the interface between the 35% and 60% Percoll were collected. The cells were washed twice in PBS prior to analysis.

### Isolation and sorting of preneoplastic lesions (PNLs)

After euthanasia, cerebella at 3-8 weeks were isolated and dissociated by trypsin followed by DNaseI treatment (*39*). The cells were resuspended in 0.1% BSA/PBS with DNaseI. The GFP-positive population representing PNLs were collected by fluorescence-activated cell sorting (FACS) using the FACS Melody system (BD Biosciences). After doublet/triplet and dead cells were excluded, GFP-positive fractions were collected for analysis. Data were analyzed using FlowJo version 10 (BD Biosciences).

### Gene expression analysis

Total RNA was extracted from P5 GCPs using RNeasy Plus Micro Kit using on-column DNA digestion protocol (Qiagen). Reverse transcription was performed using qScript (Quanta Biosciences) followed by real-time PCR using KAPA SYBR FAST ROX Low (Roche) on the ViiA 7 Real-Time PCR instrument (Applied Biosystems). The following primers were used for *Gli1* (Fwd GAG GTT GGG ATG AAG AAG CA: Rev CTT GTG GTG GAG TCA TTG GA), *Pgbd5* (Fwd GCG GCC GGA AAG AAC TAT ATC: Rev CAC AGC AGT AGA TCC CTT GC), and *Actb* (Fwd GAG AAG ATC TGG CAC CAC ACC: Rev GGT CTC AAA CAT GAT CTG GGT C).

### RNA ISH/FISH

BaseScope hybridization probes specific for *Pgbd5* exon 4 were generated as per manufacturer’s instructions (ACD, catalogue 1181898-C1). Upon cardiac perfusion and fixation in 4% PFA/0.1M PB overnight, dissected brains were washed twice with 30% sucrose/PBS and incubated overnight at 4°C. After cryopreservation in sucrose, brains were embedded in OCT (Thermo), and blocks were sectioned sagittally in 10 micrometer sections using a rotary microtome cryostat (Leica). Sections were stored at -80°C. Cryosections were baked for one hour at 60°C and fixed in 4% PFA for 15 min followed by washing in PBS. After dehydration, epitope retrieval treatment with ER2 for 5 min at 95°C and subsequent Protease III treatment for 15 min at 40°C were performed. The probe set was hybridized for two hours at 42°C. Signal amplification steps were performed according to the manufacturer’s protocol. Fast Red (Leica Bond Polymer Refine Red Detection kit DS9390) was used as chromogen. Hematoxylin was used as a counterstain. Mouse *Ppib* (ACD, catalogue 701078) and *Bacillus subtilis dapB* (ACD, catalogue 701018) probe sets were used as positive and negative controls, respectively. An adjacent section of the *Pgbd5* ISH section was used to identify *Cre*-expressing cells, i.e., tumor cells. A *Cre*-specific probe set (ACD, catalogue 312288-C2) was used for FISH. Cryosections from frozen samples were baked for one hour at 60°C. Sections were fixed in 4% PFA for 15 min and washed in PBS followed by dehydration. Epitope retrieval treatment with ER2 for 10 min at 95°C, followed by 10 min of incubation with ACD 2.5 LS Hydrogen Peroxide. The probe set was hybridized for two hours at 42°C. Mouse *Ppib* (ACD, catalogue 313918) and bacterial *dapB* (ACD, catalogue 312038) probes were used as positive and negative controls, respectively. The hybridized probes were detected using the RNAscope 2.5 LS Reagent Kit – Brown (ACD, catalogue 322100) according to manufacturer’s instructions with the following modifications. DAB application was omitted and replaced with Alexa Fluor 488 Tyramide signal amplification reagent for 20 min at room temperature (Life Technologies, B40953). After staining, slides were washed in PBS and incubated in 5 μg/ml 4’,6-diamidino-2-phenylindole (DAPI; Sigma Aldrich) in PBS for 5 min, rinsed in PBS, and mounted in Mowiol 4–88 (Calbiochem). Slides were stored at -20°C before imaging.

### Whole-genome sequencing of mouse medulloblastomas

Tissues, including tumor and matched skin or spleen were harvested from symptomatic mice and flash frozen using a dry ice and ethanol bath. DNA and RNA were extracted using the Qiagen AllPrep kit, according to manufacturer’s instructions. 2×150bp paired end libraries were prepared using Illumina TruSeq PCR free DNA kit and sequenced using Illumina HiSeq X at a depth of 80X for tumors and 40X for matched normal tissues. Reads were aligned to the mm10 reference genome with BWA-MEM (version 0.7.15) and processed by eliminating duplicate reads with NovoSort MarkDuplicates (version 3.08.02). Single nucleotide variants were detected using Mutect2 (version 4.0.5.1), Strelka2 (version 2.9.3), and Lancet (exonic) (version 1.0.7). High confidence SNVs included those that were detected by more than one caller. Structural variants were detected using Manta, SvABA (version 0.2.1), and Lumpy (version 0.2.13). For structural variants, all those that passed quality filters were included. Sequencing data are available from NCBI SRA.

### Single-nucleotide mutational signature analysis of mouse medulloblastomas

Mutational signature analysis was performed using three main steps: de novo extraction, assignment and fitting (*40*). For the first step, we ran hierarchical Dirichlet process (https://github.com/nicolaroberts/hdp). Then all extracted signatures were assigned to the COSMIC reference (https://cancer.sanger.ac.uk/signatures/sbs/) (*27*) in order to define which known mutational processes are active. Finally, we applied our recently developed fitting algorithm (https://github.com/UM-Myeloma-Genomics/mmsig) to estimate both the presence and the contribution of each mutational signature in each sample (*41*).

### Identification of genes affected by structural variants in mouse medulloblastomas

Genes analyzed included those from the COSMIC cancer gene census (*42*). We considered genes to be putatively affected by structural variants if 1) the gene was affected by a structural variant breakpoint or 2) the gene overlapped with a duplication, inversion, or deletion or 3) the gene resided within 25kb of a translocation breakpoint. For the Oncoprint depicted in Fig. 3C, we curated putative driver alterations arising from *Pgbd5*-induced SVs by analyzing for recurrence, as well as comparing to genes known to be recurrently mutated in human SHH medulloblastomas. Specifically, the putative tumor suppressor and oncogenes were identified as 1) genes affected by SVs ≥4 *Ptch1-mutant* tumors, 2) genes affected by point mutations in ≥2 tumors, 3) genes affected by SVs in ≥2 tumors that were also affected by SVs in ≥10% of human SHH MBs, or 4) genes affected by SVs that were demonstrated to be recurrently mutated in Northcott et al (*5*).

### Whole genome sequencing of human medulloblastomas

Somatic structural variants were called using Genomon-SV (v0.4.1) and DELLY2 (v0.7.5) as described in Skowron et al (*22*). For each structural variant detected by either algorithm, we used Velvet to assemble reads around the detected breakpoints, and the resultant contigs were then re-mapped locally using human genome reference sequences with and without detected structural variants using blat. This approach ensures detection of heterozygous structural variants, which would be mapped both to reference sequence without incorporated variant contigs, as well as reference sequence that incorporates them. Subsequently, we selected variants for which assembled contigs could be mapped to the reference sequence containing somatic structural variants, and variants from matched normal tissue that could not be mapped to the reference sequence that incorporates variant contig sequences. Sequencing data are available from EGA using the following accession numbers: EGAD00001003125; EGAD00001004347; EGAD00001003127.

### Identification of breakpoint sequence motifs in mouse medulloblastomas

Sequences +/- 50bp flanking both breakpoints from identified structural variants were extracted with bedtools (version 2.29.2) using the mm10 reference genome. To identify motifs that are putatively associated with Pgbd5 activity in mouse tumors, we used de novo motif analysis of the breakpoint sequences from *Pgbd5*-expressing tumors and the breakpoint sequences from *Pgbd5*-deficient tumors as background. We used discriminative MEME (v5.4.1) (*43*), which determines relative sequence enrichment in an experimental set of sequences relative to a control set. *Pgbd5*-deficient breakpoint sequences were used as controls. Discriminative MEME was used with default parameters but limited to between 11 and 16 base pairs. Repetitive motifs were eliminated, and the remainder were chosen as putative *Pgbd5*-associated motifs. For SmoM2-mutant tumors, motifs were identified using classic MEME with default parameters and limited to between 11 and 16 base pairs.

### Identification of breakpoint sequence motifs in human medulloblastomas

Sequences +/- 50bp flanking both breakpoints from identified structural variants were extracted with bedtools (version 2.29.2) using the hg19 reference genome. To identify *de novo* motifs, classic MEME (version 5.1.1) (*44*) with default parameters but limited to 11bp was used (http://meme-suite.org/). Putative PGBD5-specific signal sequence (PSS) motifs were selected by eliminating repetitive motifs and by determining whether candidate motifs were found when discriminative MEME was run relative to 50,000 randomly selected 50-mers from the hg19 reference genome without repeat masking. To quantify the abundance of motifs at structural variant breakpoints, we used FIMO (version 5.1.1) (*44*) with default parameters (Fig. S13). Q-value cutoffs for quantitation were determined by identifying a threshold where previously identified PSS motifs (Rhabdoid and RPE) could be specifically detected, but negative control motifs (RAG1/2 and scrambled) were not. This cutoff was characterized by a receiver operating characteristic (ROC) curve and chosen for sensitivity of 100% and specificity of 75% for the previously identified PSS motifs. For the quantifications, each structural variant is counted only once (i.e., each structural variant has 4×50mers and 2x breakpoints). As a comparator, we included human breast carcinoma structural variants (*3, 23*). To mimic the background distribution of the FIMO algorithm, we employed randomly scrambled sequences.

### Sequence motif comparisons

The similarity of motifs was evaluated using TomTom (https://meme-suite.org/meme/tools/tomtom) (*45*). The first set of query motifs which included those from 3 *Ptch1*-mutant tumors and 6 *SmoM2*-mutant tumors were compared with previously identified target motifs (PSS motifs: including 11 motifs (PSS_1 to 11), PSS_Rhab, PSS_RPE, and PSS_HPRT). To test for similarities, 10,000 shuffled target motifs were generated and their p-values were examined. If the p-value of the target motif ranked within the lowest 5% of all 10,001 p-values, motif was deemed significantly similar. In the same way, the second set of query motifs (hMB1-4) were compared with nine mouse motifs and their shuffled sequences. Finally, the query motif sets (hMB1-4) were compared with the previously identified 14 PSS motifs and their shuffled sequences.

---

**Fig. S1.**
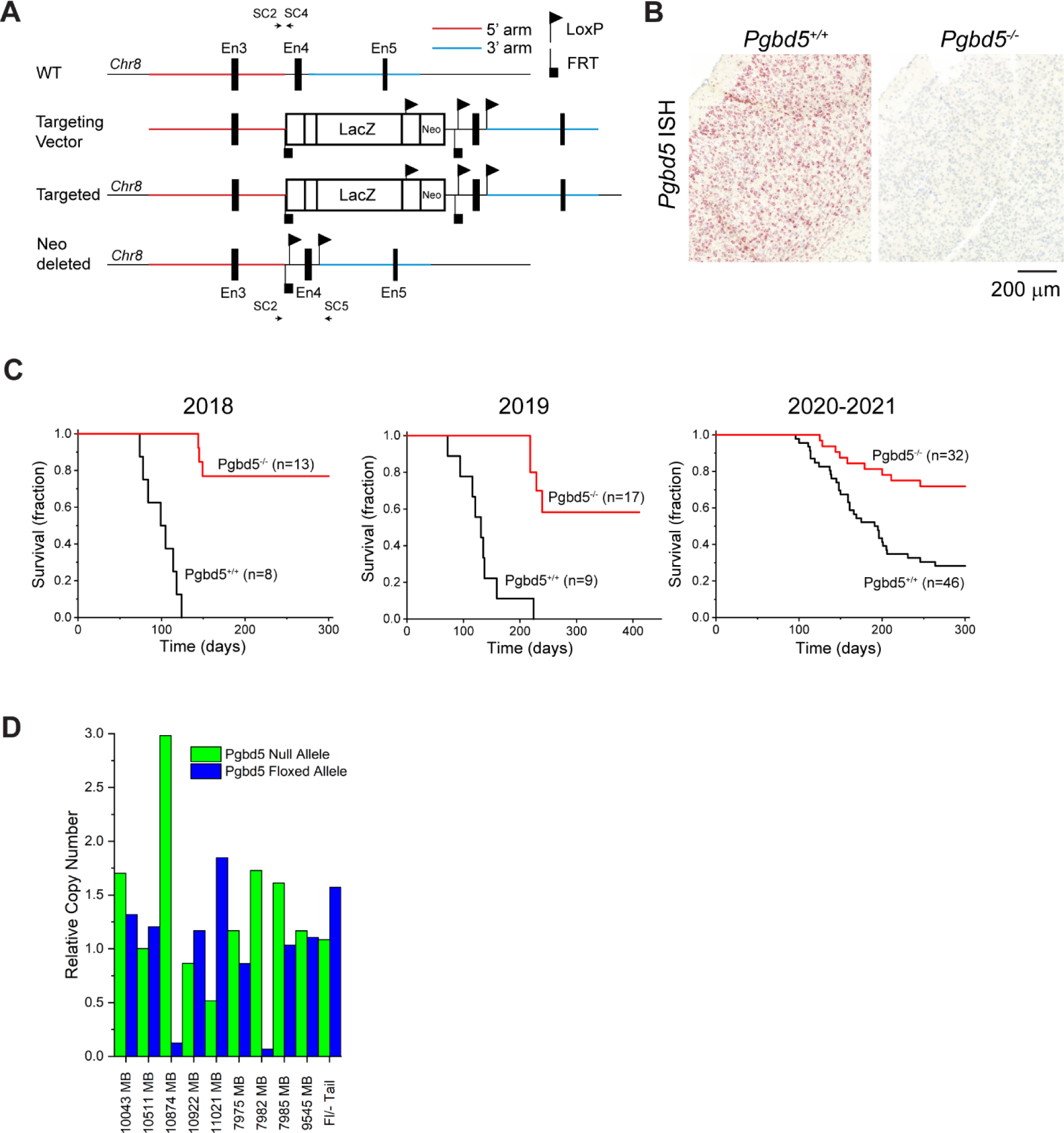
(**A**) Generation of *Pgbd5^-/-^* mice. The top shows the *Pgbd5* wildtype (WT) gene locus including exon (En) 3-5 at chromosome 8. The targeted vector (second from top) consists of LacZ and neoR cassettes flanked by FRT sequences (black inverted square flag) and exon 4 flanked by LoxP sequences (black inverted triangle flag). Red and blue regions are 5’ and 3’ homology arms, respectively. The third line shows a successfully targeted gene locus which is confirmed by genotyping using SC2 and SC4 primers. These successfully targeted ES cell clones were microinjected into Balb/c blastocysts and resulting chimeras were crossed with FLP deleter mice to remove the LacZ and neoR cassettes. The bottom line shows the Pgbd5 floxed allele after removal of the LacZ/neoR cassettes. This individual was crossed with an EIIa-Cre mice to generate global *Pgbd5*^-/-^ mice, as confirmed by genotyping with SC2 and SC5 primers. (**B**) Representative images of the ISH using a BaseScope probe set against exon 4 of *Pgbd5* transcripts. Sagittal sections of *Pgbd5^+/+^* (left) and *Pgbd5^-/-^*(right) cortices show *Pgbd5^+/+^*-specific signal in red. Nuclei were counter-stained with Hematoxylin (blue). (**C**) Three independent survival analyses in the *Ptch1*-mutant model conducted in 2018, 2019, and 2020-2021. *Pgbd5^-/-^* (red) mice exhibited significant protection from tumor development in all three cohorts (log-rank *p* = 3.0e-7, 1.0e-7, and 1.9e-4). (**D**) Allele copy number analysis of nine *Pgbd5^fl/-^* tumor and one tail tissues by genomic qPCR. Green and blue indicates allele copy numbers of null and floxed alleles relative to those of *jun*, respectively. 10874 and 7982 tumors lost the floxed alleles while the other seven tumors retain the floxed alleles.

**Fig. S2.**
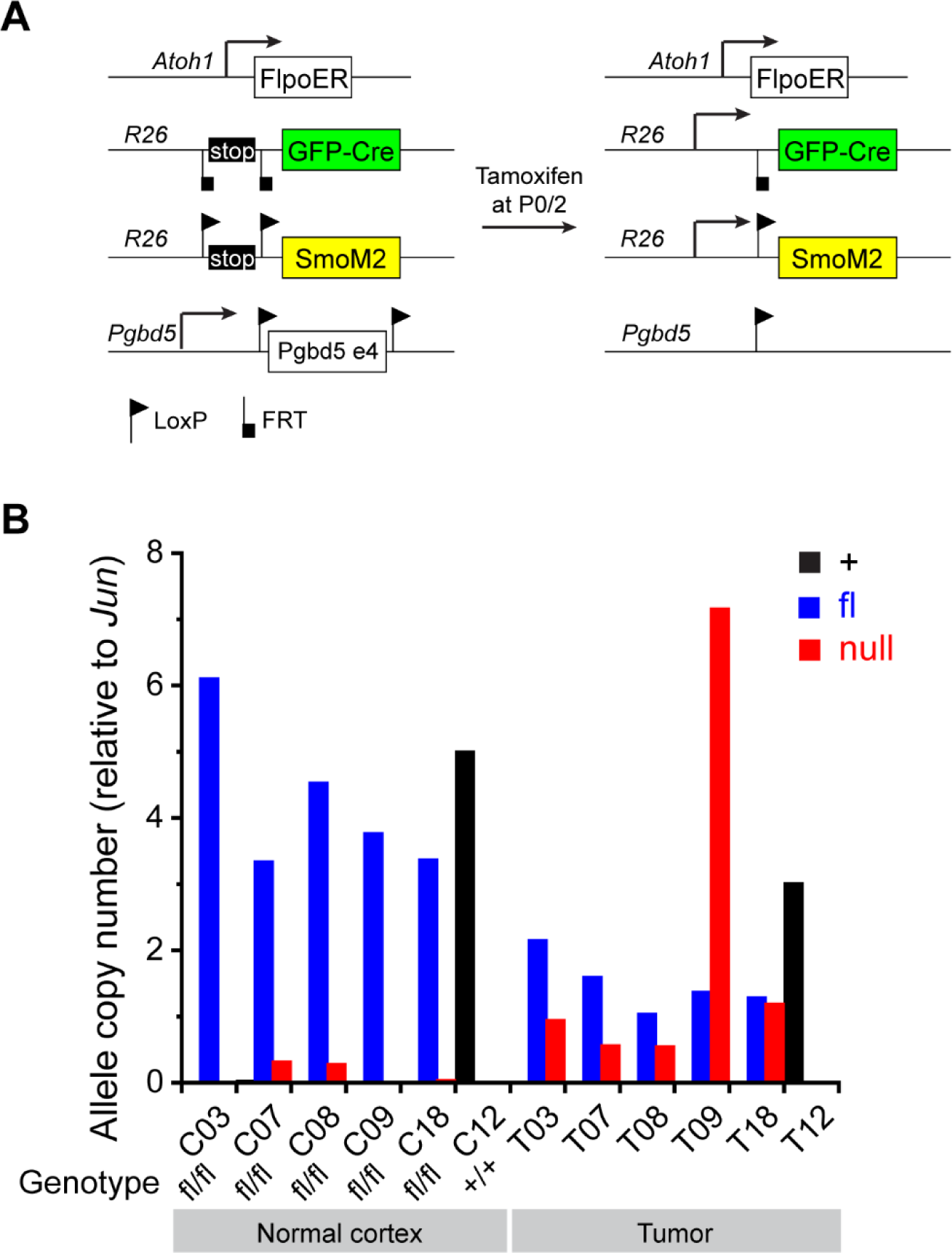
(**A**) Schematic showing strategy of the MASTR approach. Upon Tamoxifen injection at P0, FlpoER removes a stop cassette to drive GFP-Cre (green) expression, which induces SmoM2 protein (yellow) expression and concurrent deletion of *Pgbd5*. Triangle and square flags indicate LoxP and FRT sequences, respectively. (**B**) Allele copy number analysis of five normal cortices (left) and their corresponding tumors (right) by genomic qPCR. C03, C07, C08, C09, and C18 are *Pgbd5^fl/fl^*; C12 is *Pgbd5^+/+^*. Black, blue, and red indicate wildtype, floxed, and null alleles, respectively. In the tumors (T), except for T09, the floxed alleles are retained.

**Fig. S3.**
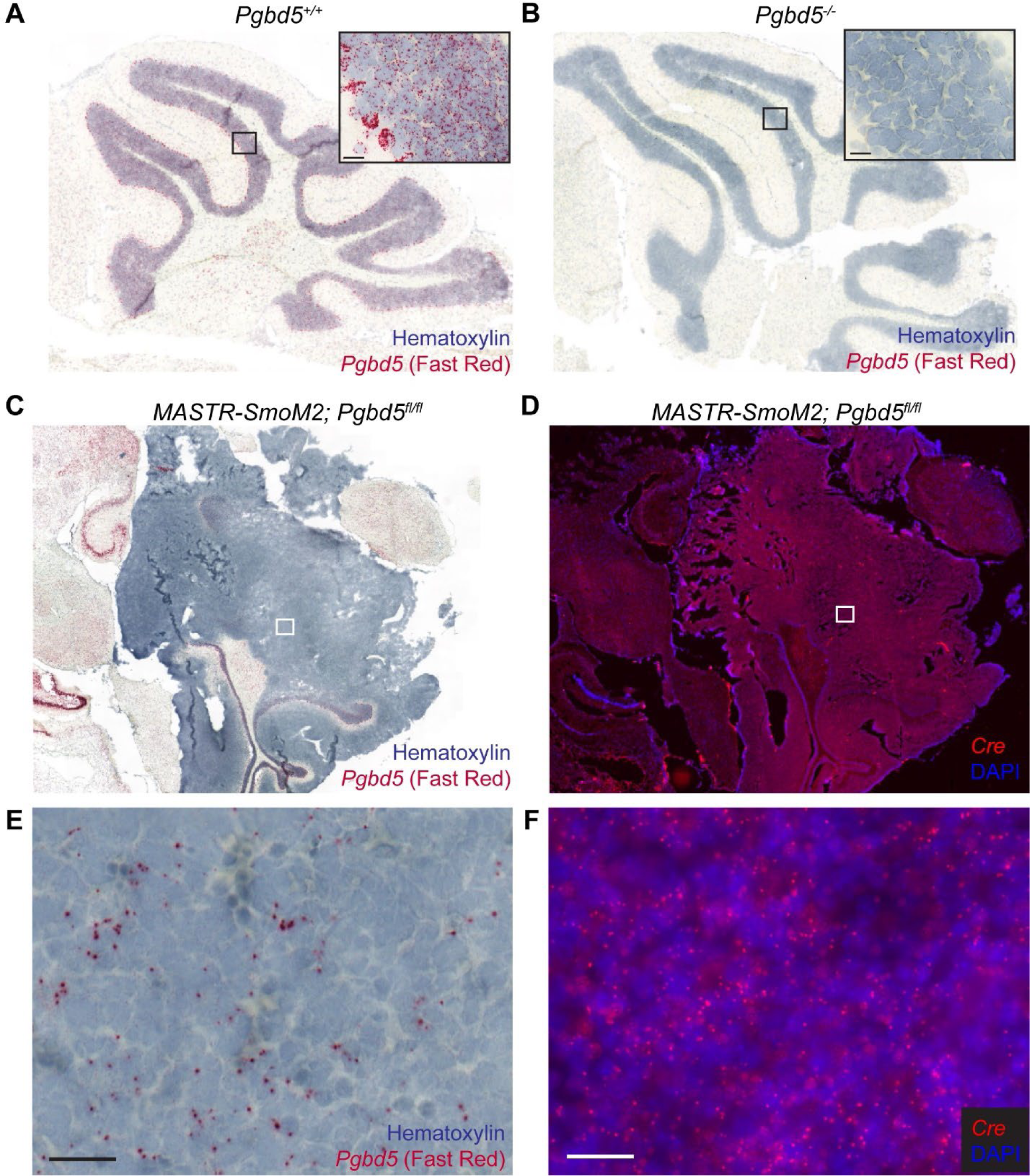
*In situ* hybridization (ISH) images of BaseScope *Pgbd5* probe set for *Pgbd5^+/+^* (positive control) (**A**) and *Pgbd5^-/-^* (negative control) (**B**) cerebella. *Pgbd5* signal is detected by Fast Red in *Pgbd5^+/+^* cerebellum whereas no signal is observed in *Pgbd5^-/-^*. Nuclei are counterstained with hematoxylin. High magnification images are shown in the black insets. (**C**) Representative *Pgbd5* ISH image of a tumor from the *MASTR-SmoM2; Pgbd5^fl/fl^* mouse. (**D**) *Cre* FISH image of the adjoining section of (**C**). Red fluorescence represents the *Cre* signal. Nuclei are counterstained with DAPI. (**E**) and (**F**) are higher magnification images of tumor areas (white rectangles) in (**C**) and (**D**), respectively. Residual *Pgbd5* signal is observed in the tumor (**E**) which is marked by the homogeneous *Cre* signal (**F**). Scale bars: 20 μm.

**Fig. S4.**
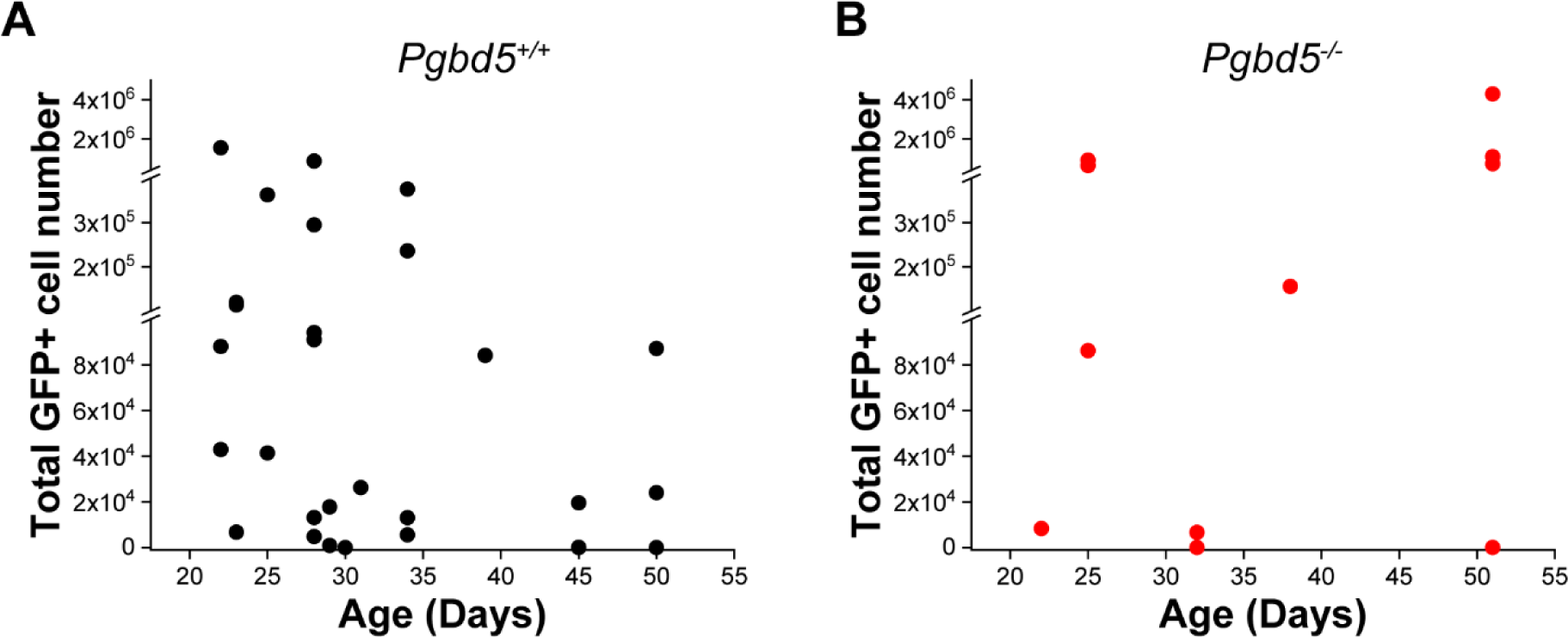
Total Atoh1-GFP+ cell numbers in the *Pgbd5^+/+^* (**A**) and *Pgbd5^-/-^* (**B**) cerebella between 3 and 8 weeks by FACS. Both show similar numbers of the total Atoh1-GFP+ cells (t-test *p* = 0.16).

**Fig. S5.**
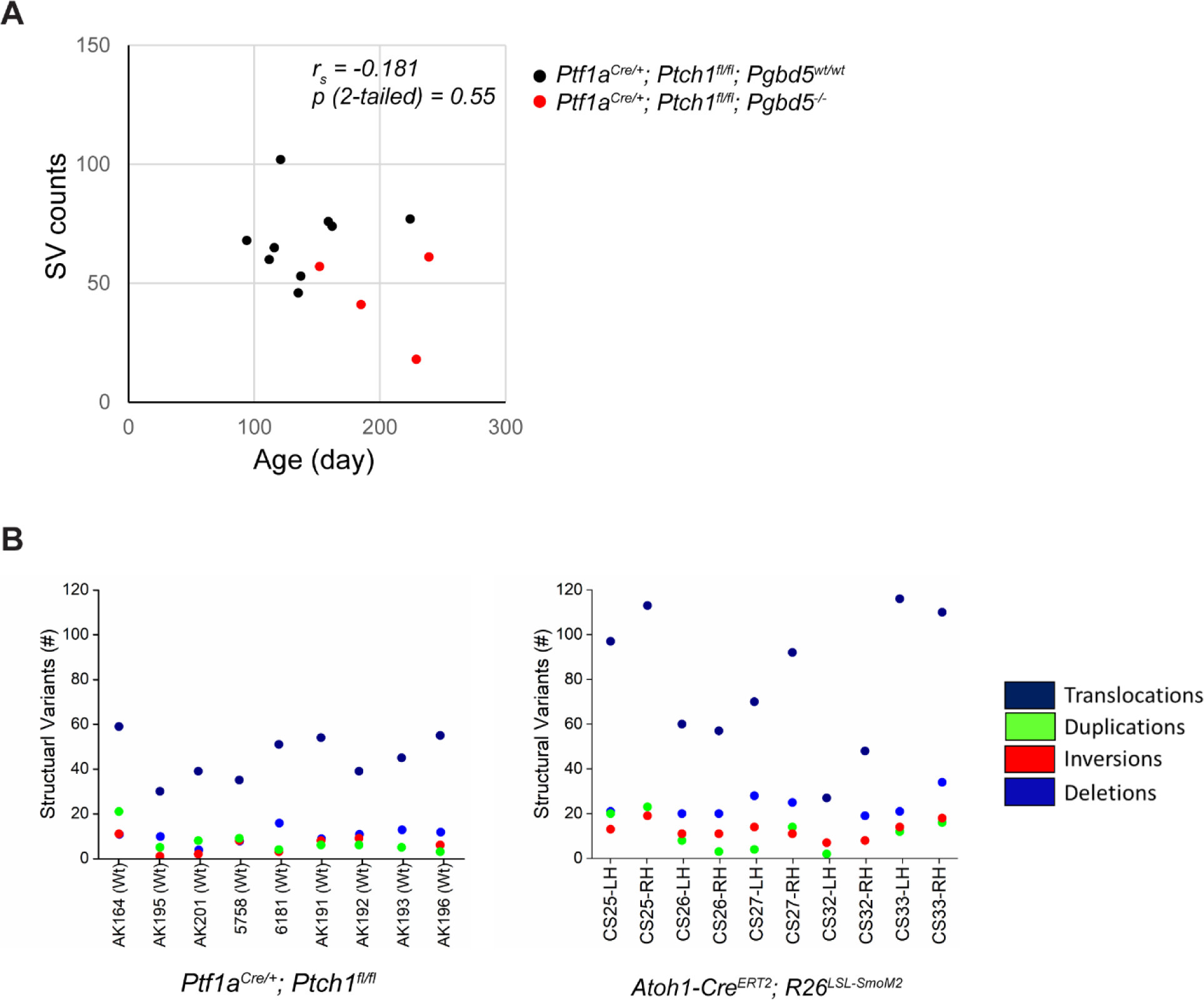
(**A**) Scatter plot showing the relationship between the age of tumors and numbers of somatic structural variants in the *Ptch1*-mutant model. Both *Pgbd5^+/+^* (black) and *Pgbd5^-/-^* (red) tumors did not show significant correlation between age and numbers of structural variants. (**B**) Numbers of different types of structural variants in *Ptch1-* (left) and *SmoM2*-mutant (right) tumors. Both models show similar numbers of mutation types.

**Fig. S6.**
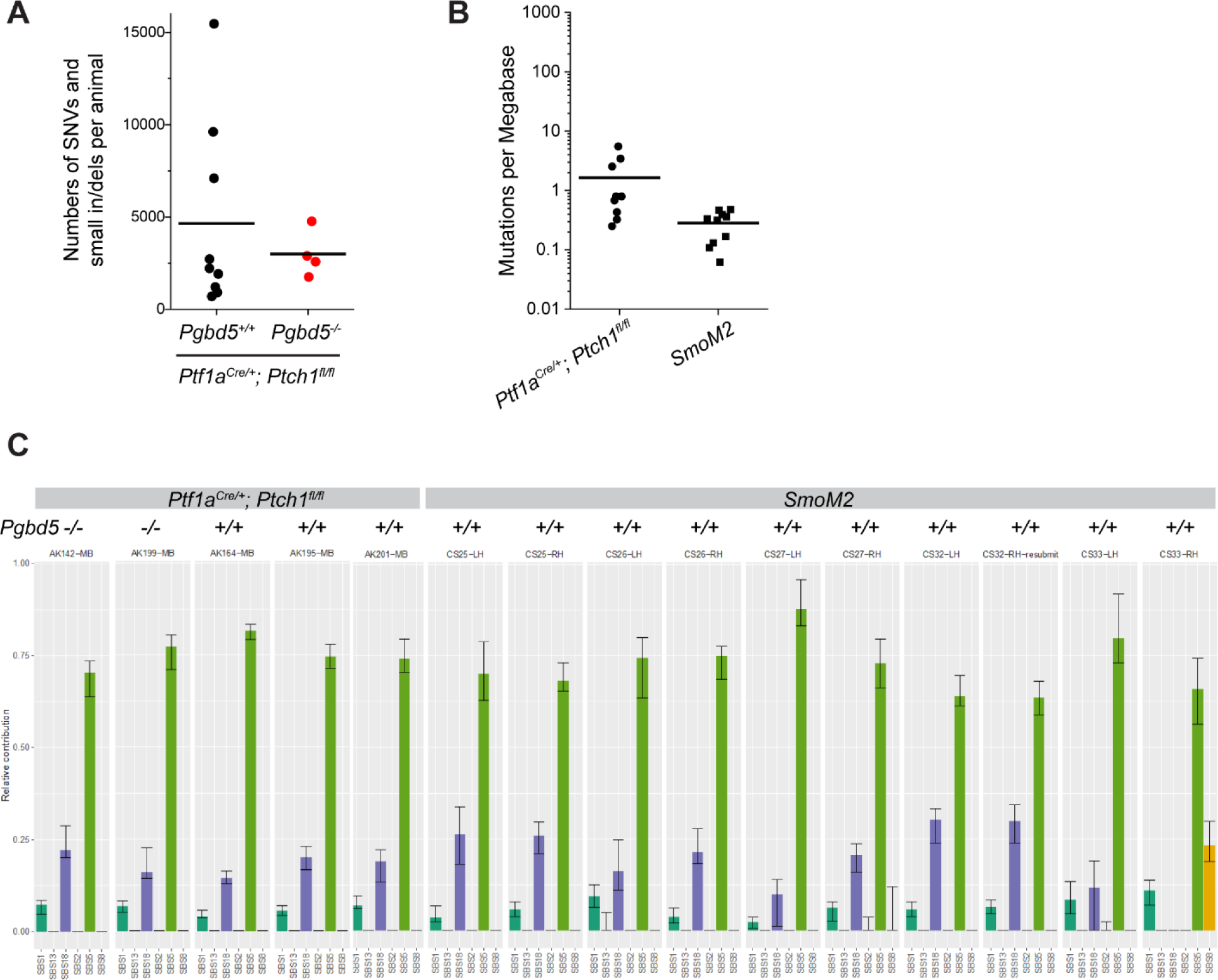
(**A**) Comparison of numbers of SNVs and small indels between *Pgbd5^+/+^* (black: N = 9) and *Pgbd5^-/-^* (red: N = 4) *Ptch1*-mutant tumors. Both tumors have similar numbers of SNVs and small indels. Bars indicate means. (**B**) Comparison of numbers of SNVs and small indels between *Ptch1*- (N = 9) and *SmoM2*- (N = 10) mutant tumors. Both tumors show relatively low numbers of mutations. (**C**) Mutational signature analysis. Ratios of contribution of the major signatures found in SNVs. The left five are from the *Ptch1*-mutant tumors (2 *Pgbd5^-/-^* and 3 *Pgbd5^+/+^*) and the right ten are from the *SmoM2*-mutant tumors. SBS1; dark green, SBS5; light green, SBS18; purple, SBS8; orange. SBS5 is the most dominant signature, followed by SBS18 and SBS1. The only exception is *SmoM2*-mutant CS33-RH exhibiting the SBS8 contribution instead of SBS18.

**Fig. S7.**
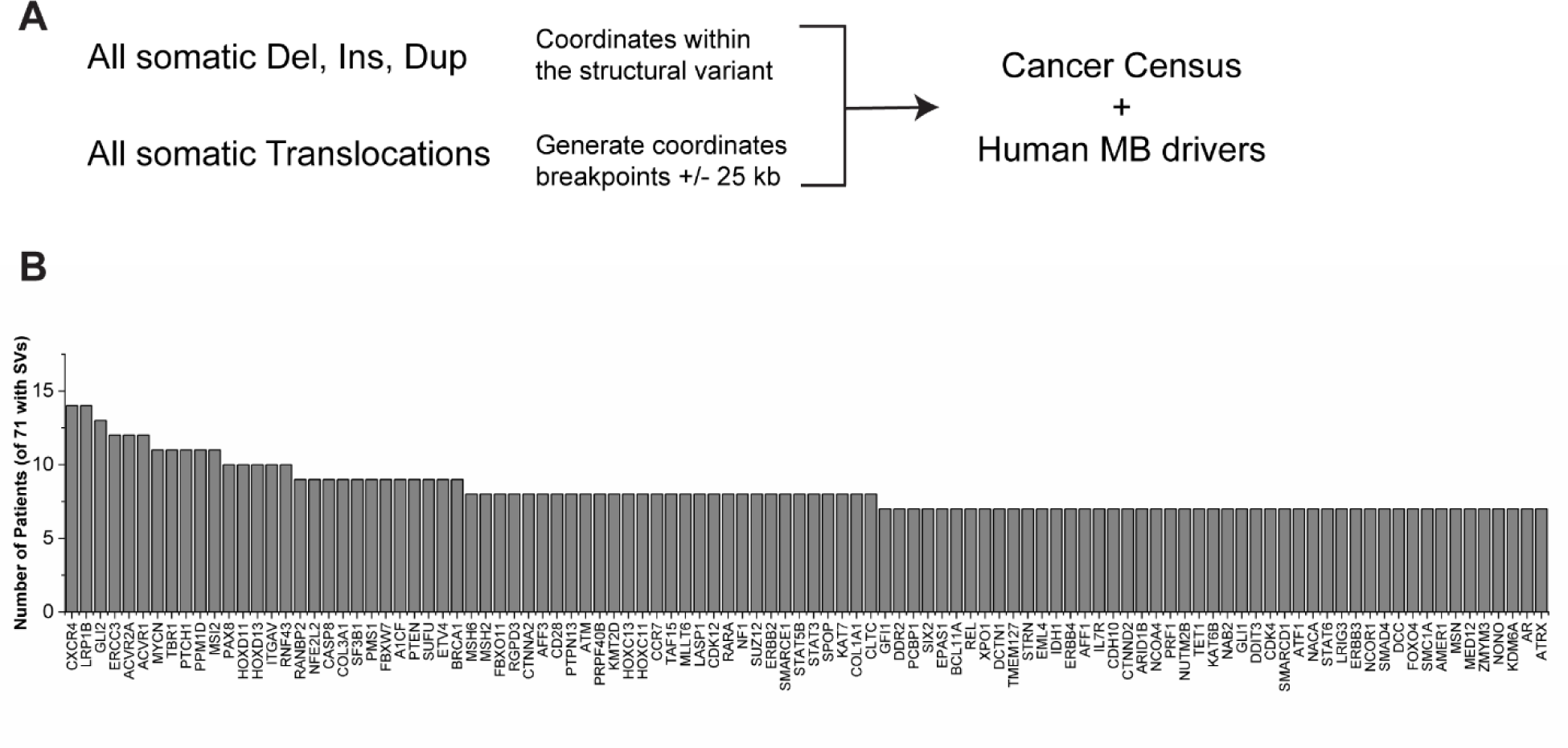
Cancer census genes affected by somatic structural variants in a cohort of human SHH medulloblastomas. (**A**) Schematic to identify genes putatively affected by SVs. Genes were considered affected by structural variants if the gene was intersected by a structural variant breakpoint, if the gene was between a duplication, inversion, or deletion, or if the gene was within 25kb of a translocation breakpoint. (**B**) Many recurrent medulloblastoma tumor suppressors and oncogenes are affected by SVs. In the cohort of SHH subtype medulloblastoma (Fig. 4A), 71 tumors had structural variants. Genes known to be recurrently affected in human medulloblastomas are affected in >10% of cases, including *CXCR4, MYCN, PPM1D, PTCH1, GLI2*, and others.

**Fig. S8.**
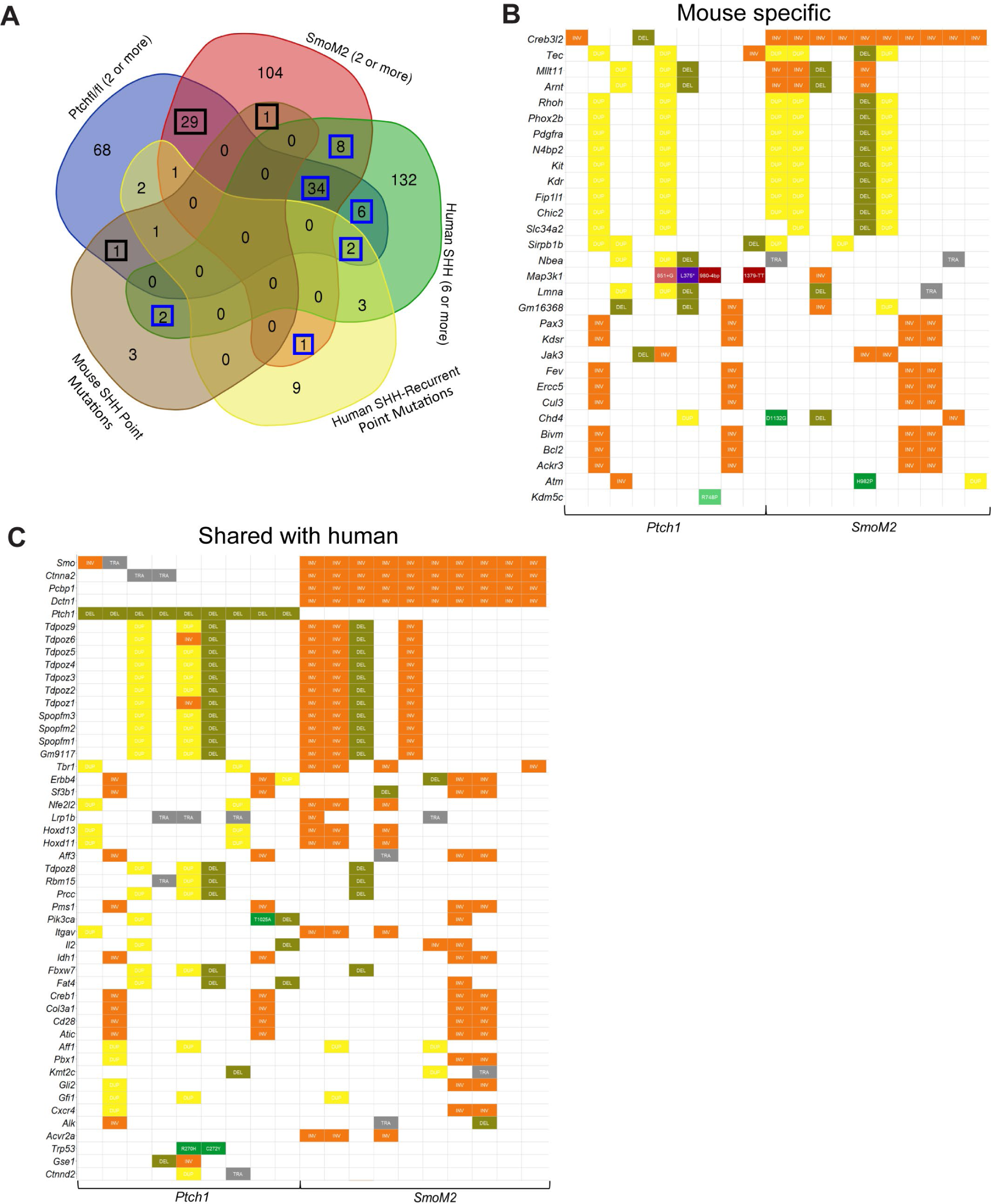
Genes recurrently affected by somatic structural variants in human medulloblastoma are also affected in *Pgbd5^+/+^; Ptch^fl/fl^* medulloblastomas and *SmoM2*-mutant medulloblastoma. (**A**) Venn diagram depicting the overlap among cancer census genes between human medulloblastomas and each mouse tumor model, including those affected by structural variants and point mutations. The numbers boxed in black are mouse specific genes (B), and the numbers boxed in blue represent overlaps between human and mouse tumors (C). (**B**) Genes affected in two or more mouse tumors that are not affected by structural variants in more than 10% of SHH medulloblastomas in this cohort. Affected genes are chosen with the same criteria as in Fig. S7A. (**C**) Set of genes affected in both human and mouse medulloblastomas, including structural variants and point mutations.

**Fig. S9.**
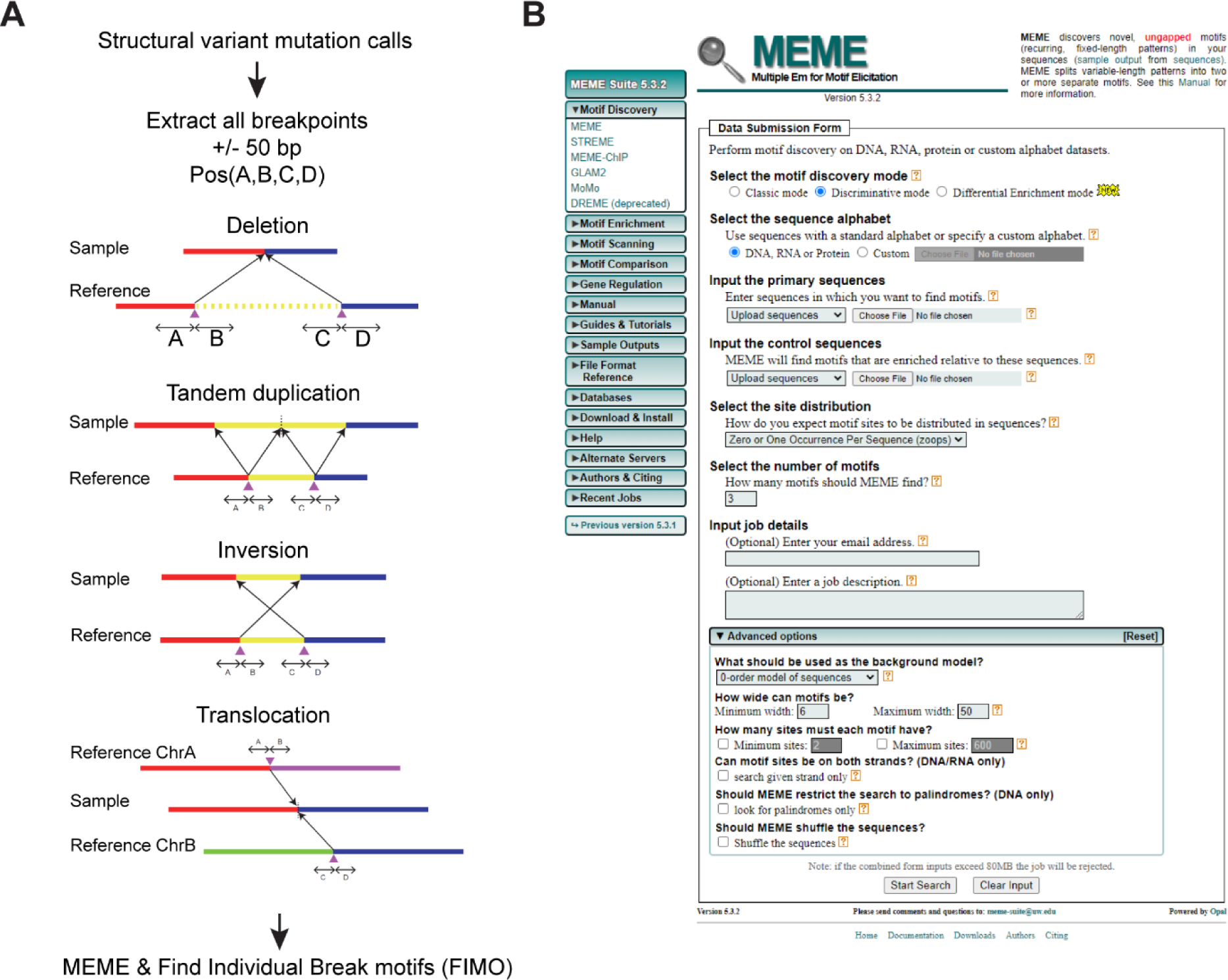
**(A)** Schematic describing the method used to extract 50mer flanking sequences from structural variant breakpoints in mouse medulloblastoma genomes. Following somatic mutation detection, 50mer sequences were extracted using bedtools in FASTA format compatible with MEME (see methods). (**B**) Depiction of paremters for MEME analysis. Discriminative MEME was used with default settings. *Pgbd5*-wildtype tumor 50mers were used as the primary sequences (2480 50mers) and *Pgbd5^-/-^* tumor 50mers were used as the control sequences (708 50mers). MEME was used with 11, 12, 13, 14, 15, and 16 maximum base pairs. Repetitive sequences were eliminated and specific sequences were chosen as putative *Pgbd5* specific motifs.

**Fig. S10.**
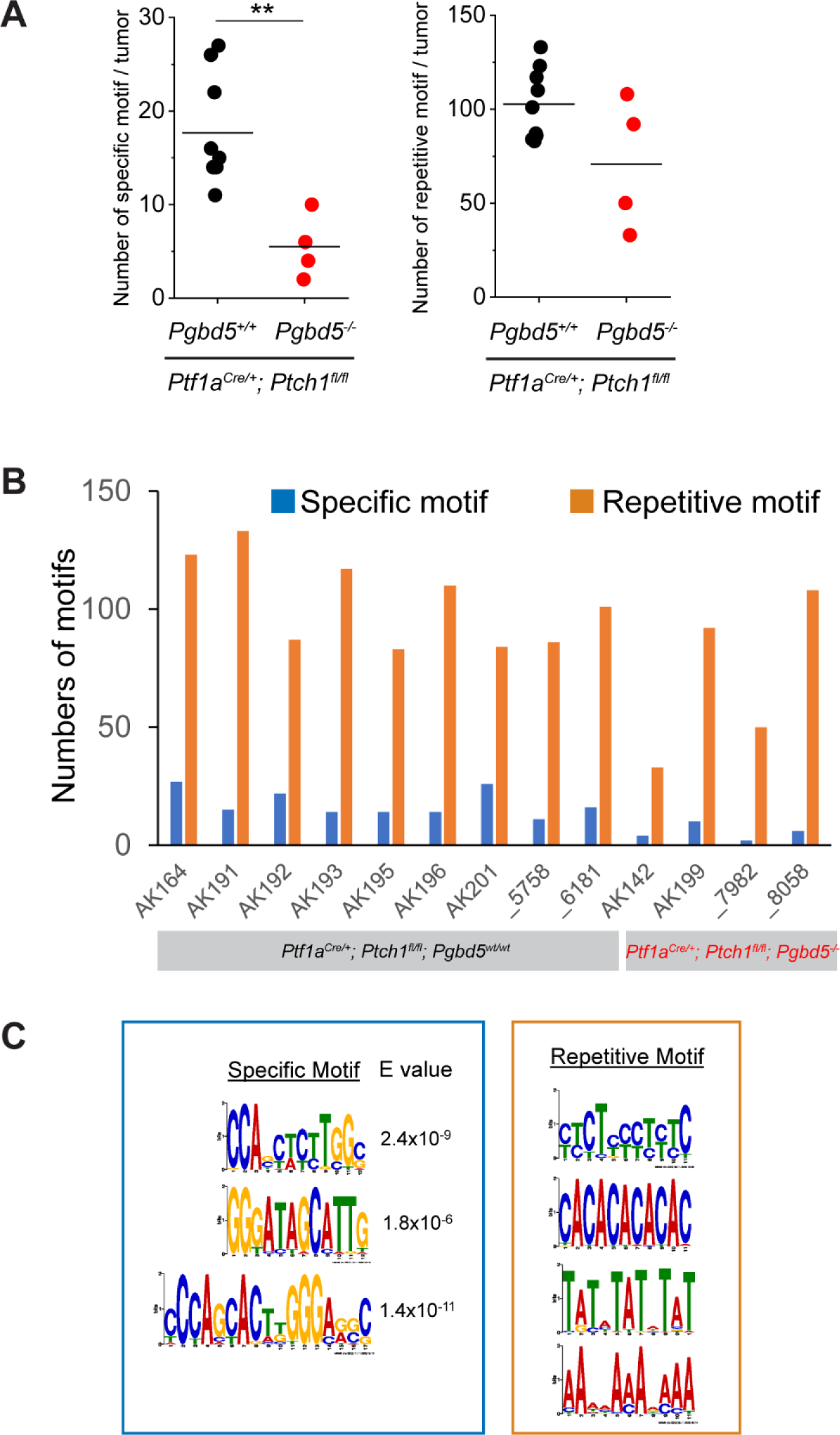
**(A)** Comparison of the number of Pgbd5-specific motifs shown in the left diagram of (C) between *Pgbd5^+/+^* (black) and *Pgbd5^-/-^*(red) *Ptch1*-mutant tumors (Left graph). *Pgbd5^+/+^* tumors harbor more specific motifs (t-test *p* = 2.7e-3). Right graph shows comparison of numbers of the repetitive motifs shown in the right diagram of (C) between *Pgbd5^+/+^* and *Pgbd5^-/-^* tumors. No significant difference in the number of repetitive motifs was observed. **(B)** Numbers of motifs in each tumor from the *Ptch1*-mutant model. Left nine and right four tumors are from *Pgbd5^+/+^* and *Pgbd5^-/-^* mice, respectively. Blue indicates the total numbers of the three Pgbd5-specific motifs depicted in the left of (C) and orange indicates total numbers of the repetitive motifs shown in the right diagram of (C).

**Fig. S11.**
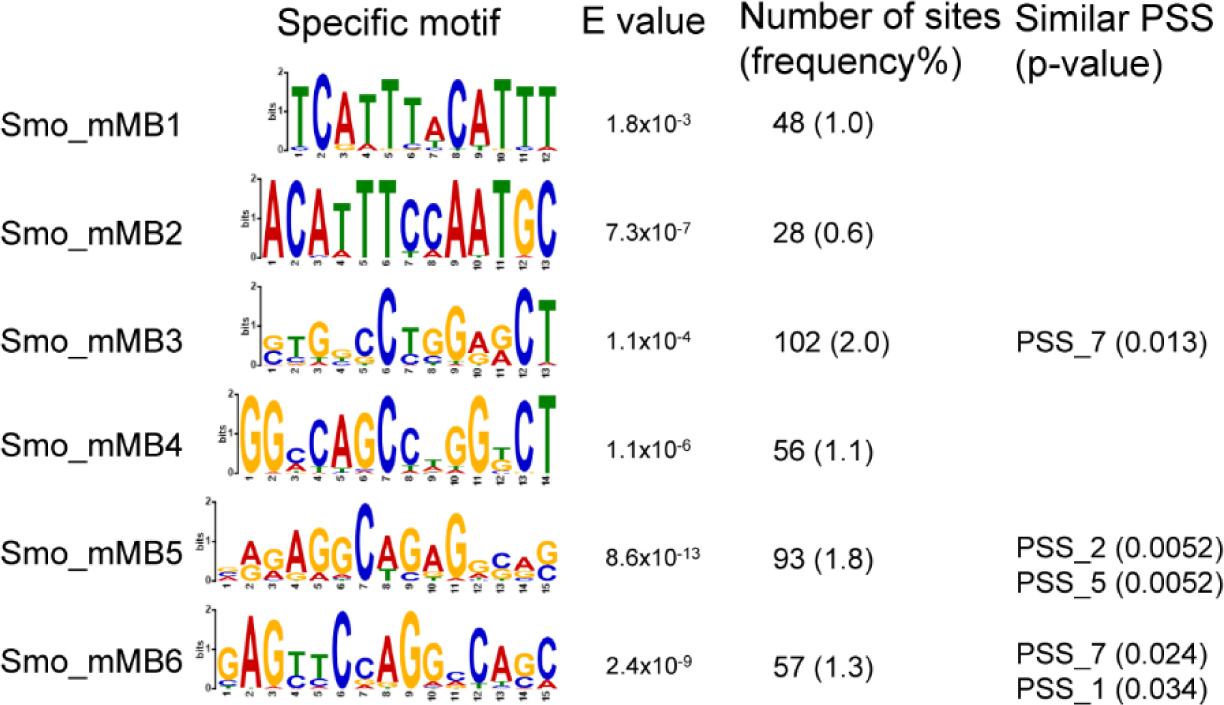
Specific motifs identified in *SmoM2*-mutant tumors share similarity with previously identified PSS motifs. E-value indicates significance based on the number of motifs that are expected should the motifs be shuffled. Frequency is the instance number within the set of breakpoint 50mers. Some of these motifs were similar to previously identified PSS motifs, as measured using TomTom.

**Fig. S12.**
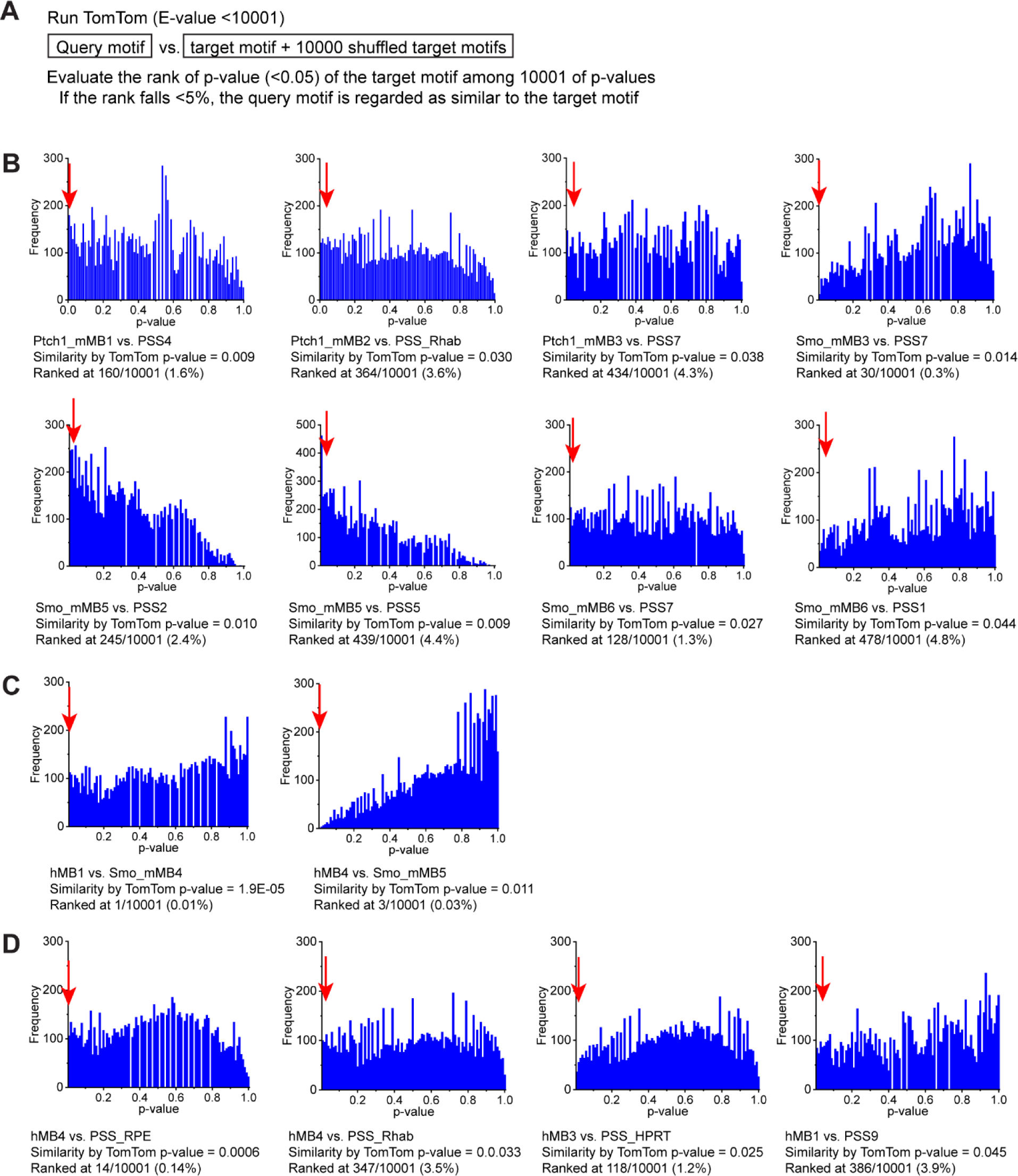
Verification of similarities among breakpoint motifs. (**A**) Method description on how the similarity is verified. The shuffled target motifs were used as background. If the *p*-value of a target motif falls within the lowest 5% of all p-values, the similarity is verified. (**B**) Histograms of all the *p*-values (target motif plus its 10,000 shuffled motifs) for mouse medulloblastoma (mMB) motifs vs. previously identified PSS motifs. Red arrows point to the *p*-value of target motifs. The *p*-value and its ranking are shown under the histogram. (**C**) Comparison of human medulloblastoma (hMB) motifs vs. mMB motifs. (**D**) Comparisons of hMB motifs vs. previously identified PSS motifs.

**Fig. S13.**
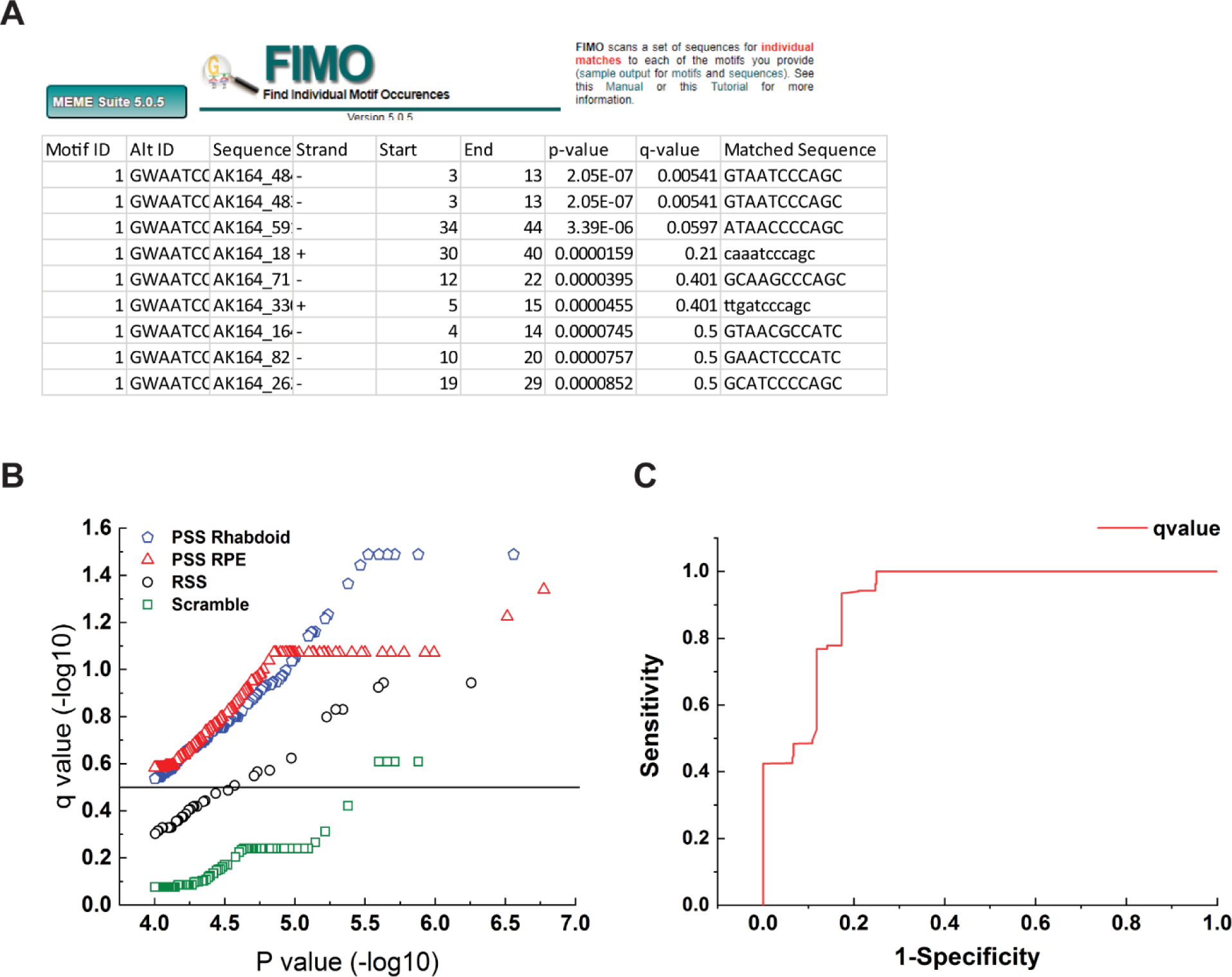
**(A)** An example of FIMO analysis. The *q*-value is a modification of the *p*-value, corrected for the respective false discovery rate. Each row represents a sequence that matched to the queried motif. **(B)** Graphical representation of the output when 50mers from human SHH subtype structural variants were queried with different motifs, including previously identified PSS motifs (Rhabdoid and RPE) and motifs known not to be associated with Pgbd5 activity, including the RAG1/2 specific signal sequence and a scrambled motif. The line indicates the *q*-value above which all sequences with PSS sequences are correctly identified, with negligible number of negative control sequences. This threshold was chosen as the q-value for breakpoint sequence motif identification. **(C)** Receiver operating characteristic curve analysis of values from (B), demonstrating the separation of PSS sequences from negative controls with 100% sensitivity and 75% specificity using q-value cutoff of 0.3.

**Fig. S14.**
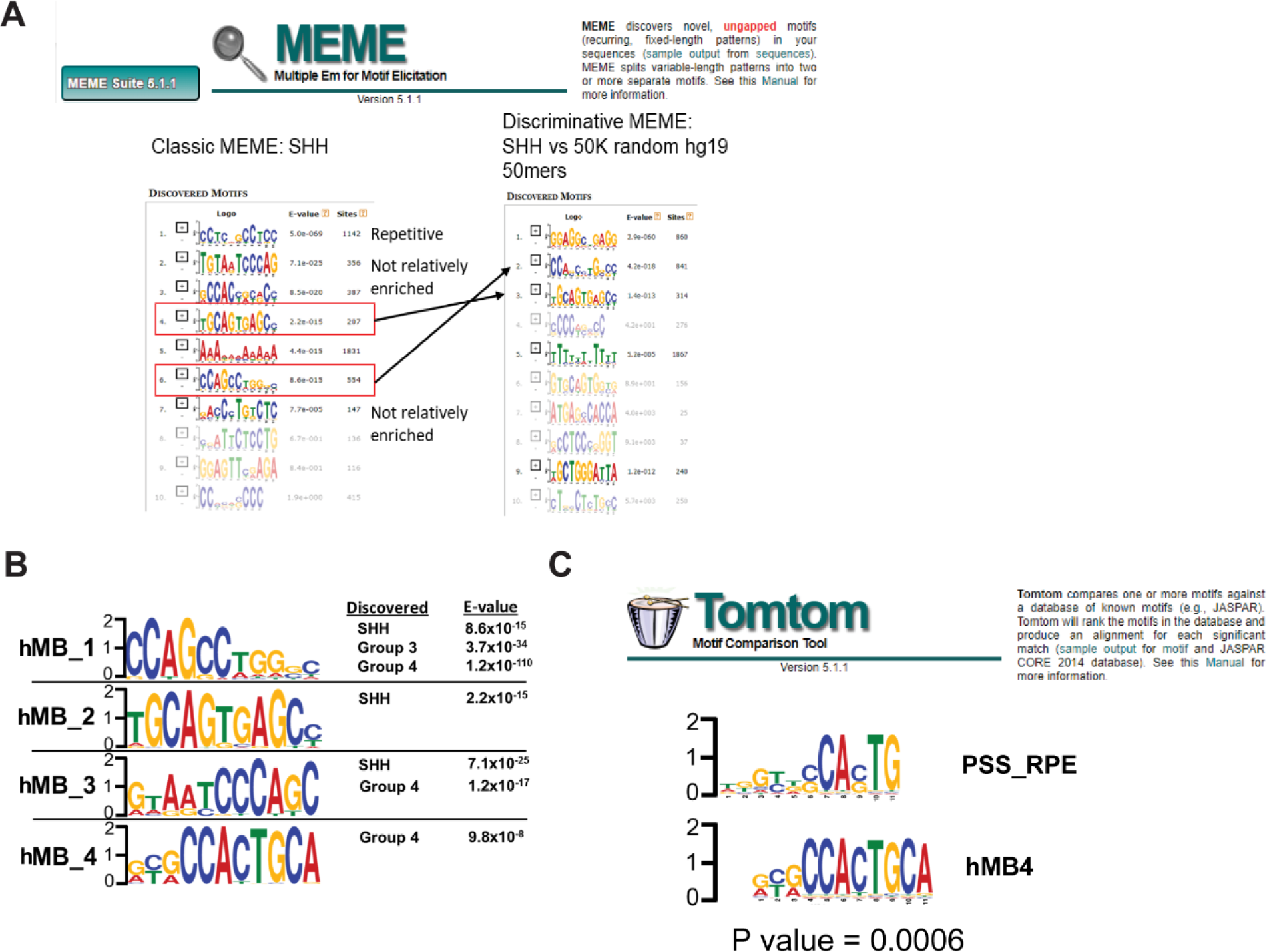
Specific sequence motifs enriched in human medulloblastomas at structural variant breakpoints. (**A**) Example output from MEME and method for eliminating repetitive motifs. Motifs were chosen based on whether they were sequence-specific (i.e., not repetitive) and whether they were enriched using discriminative MEME relative to a set of 50,000 50mers randomly sampled from the hg19 reference genome. (**B**) Specific motifs identified in human medulloblastomas. The “discovered” column describes the subtype of medulloblastoma where the motif was initially discovered based on the method in A. The *E*-values list strengths of apparent associations. (**C**) Example of motif comparison between a de novo discovered motif (hMB_4) and a previously identified Pgbd5-specific signal sequence motif (PSS_RPE).

**Table S1.**
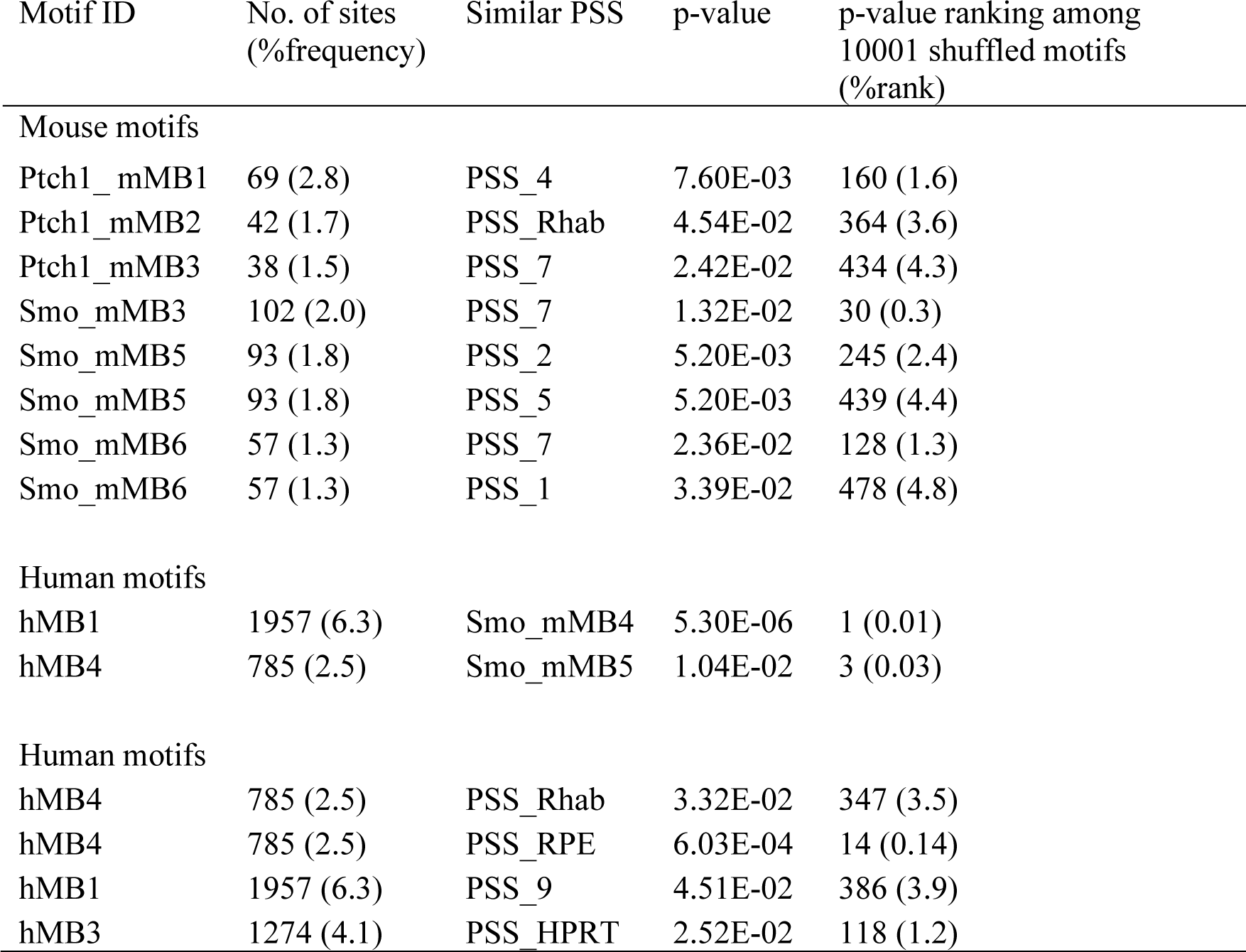
Combinations of similar motifs by motif comparison.

| Motif ID | No. of sites<br>(%frequency) | Similar PSS | p-value | p-value ranking among<br>10001 shuffled motifs<br>(%rank) |
| --- | --- | --- | --- | --- |
| Mouse motifs |  |  |  |  |
| Ptch1_mMB1 | 69 (2.8) | PSS_4 | 7.60E-03 | 160 (1.6) |
| Ptch1_mMB2 | 42 (1.7) | PSS_Rhab | 4.54E-02 | 364 (3.6) |
| Ptch1_mMB3 | 38 (1.5) | PSS_7 | 2.42E-02 | 434 (4.3) |
| Smo_mMB3 | 102 (2.0) | PSS_7 | 1.32E-02 | 30 (0.3) |
| Smo_mMB5 | 93 (1.8) | PSS_2 | 5.20E-03 | 245 (2.4) |
| Smo_mMB5 | 93 (1.8) | PSS_5 | 5.20E-03 | 439 (4.4) |
| Smo_mMB6 | 57 (1.3) | PSS_7 | 2.36E-02 | 128 (1.3) |
| Smo_mMB6 | 57 (1.3) | PSS_1 | 3.39E-02 | 478 (4.8) |
| Human motifs |  |  |  |  |
| hMB1 | 1957 (6.3) | Smo_mMB4 | 5.30E-06 | 1 (0.01) |
| hMB4 | 785 (2.5) | Smo_mMB5 | 1.04E-02 | 3 (0.03) |
| Human motifs |  |  |  |  |
| hMB4 | 785 (2.5) | PSS_Rhab | 3.32E-02 | 347 (3.5) |
| hMB4 | 785 (2.5) | PSS_RPE | 6.03E-04 | 14 (0.14) |
| hMB1 | 1957 (6.3) | PSS_9 | 4.51E-02 | 386 (3.9) |
| hMB3 | 1274 (4.1) | PSS_HPRT | 2.52E-02 | 118 (1.2) |

**Data S1. (separate file)**

SNVs and indels from WGS of Ptch1-mutant mouse MBs

**Data S2. (separate file)**

SVs from WGS of Ptch1-mutant mouse MBs

**Data S3. (separate file)**

SNVs and indels from WGS of SmoM2-mutant mouse MBs

**Data S4. (separate file)**

SVs from WGS of SmoM2-mutant mouse MBs

**Data S5. (separate file)**

Cancer census genes affected by SVs in mouse MBs

**Data S6. (separate file)**

50mer sequences extracted from mouse WGS

**Data S7. (separate file)**

MEME results with the 50mer seqs of SVs from Ptch1-mutant Pgbd5+/+ and Pgbd5-/- mice

**Data S8. (separate file)**

Cancer census genes affected by SVs in human SHH-MBs

**Data S9. (separate file)**

50mer sequences of SVs from human SHH-MBs

**Data S10. (separate file)**

50mer sequences of SVs from human Group3-MBs

**Data S11. (separate file)**

50mer sequences of SVs from human Group4-MBs

**Data S12. (separate file)**

50mer sequences of SVs from human WNT-MBs

**Data S13. (separate file)**

50mer sequences of SVs from human breast carcinoma

**Data S14. (separate file)**

FIMO output from human SHH-MBs

**Data S15. (separate file)**

FIMO output from human Group3-MBs

**Data S16. (separate file)**

FIMO output from human Group4-MBs

**Data S17. (separate file)**

FIMO output from human WNT-MBs

**Data S18. (separate file)**

FIMO output from human breast carcinoma

**Data S19. (separate file)**

SV IDs and corresponding 50mers for select MB oncogenes and suppressors

